# KGRACDA: A Model Based on Knowledge Graph from Recursion and Attention Aggregation for CircRNA-disease Association Prediction

**DOI:** 10.1101/2023.12.04.569883

**Authors:** Ying Wang, Maoyuan Ma, Yanxin Xie, Qinke Peng, Hongqiang Lyu, Hequan Sun, Laiyi Fu

**Author notes:** These authors contributed to the work equllly (Ying Wang, Maoyuan Ma).

## Abstract

CircRNA has been found to exhibit with human diseases, rendering the investigation of the circRNA-disease association (CDA) of significant importance. Deep learning provides a novel avenue for predicting CDA, however, due to the limited availability of comprehensive CDA datasets, most existing methods operate by directly reasoning over triplets of embedded entities and relations, neglecting the explicit capture local depth information. In this study, we propose a novel approach called Knowledge Graph-Enhanced Recursive Aggregation for CDA (KGRACDA), which enables the querying of explicit local structures surrounding triplets, thereby serving as crucial evidence for predicting CDA. First, we construct a comprehensive CDA knowledge graph utilizing a large-scale, multi-source heterogeneous dataset. Subsequently, we develop a graph neural network using recursive techniques and employ attention aggregation to generate CDA prediction scores. Experimental results demonstrate that KGRACDA surpasses other state-of-the-art methods by explicitly capturing local depth information related to CDA. To improve the convenience of CDA prediction, we have updated an interactive web platform named HNRBase v2.0, which provides rich visualization of circRNA data, and allows users to easily download data and predict CDA using KGRACDA. The code and datasets are publicly available at https://github.com/maoyuanma/KGRACDA.

## INTRODUCTION

Circular RNA (CircRNA), a distinctive RNA species, was initially identified by researchers back in 1976. Unlike other RNAs, circRNA exhibits a covalently closed-loop structure, wherein its 3’ and 5’ ends are intricately linked to one another^1^. Consequently, circRNA possesses the remarkable trait of resisting degradation by exonucleases, endowing it with greater stability in comparison to the majority of linear RNAs. In recent years, circRNA has been ubiquitously discovered across eukaryotic organisms, assuming a pivotal role in a myriad of physiological and pathological processes, with particular prominence in human diseases such as diabetes, cancer, and cardiovascular diseases^2^.

Therefore, studying the relationship between circRNA and diseases can provide new ideas and methods for the research of disease pathogenesis, diagnosis and treatment, which is of great significance for the prevention and treatment of diseases. However, traditional biological methods represented by high-throughput sequencing technology have high requirements for experimental equipment, complex experimental operations, and low accuracy, which restrict the research of circRNA-disease relationship^3^.

To solve this problem, a series of computational methods have been proposed in recent years to identify potential circRNA-disease associations (CDA). These methods can be roughly divided into three categories, including methods based on classical machine learning, methods based on network links, and methods based on deep learning. First, traditional machine learning methods transform the problem of predicting circRNA-disease relationships into a classification or regression problem, using different feature extraction methods and machine learning algorithms to train and test the models. For example, Peng et al.^4^ combined robust non-negative matrix factorization and label propagation (RNMFLP) to infer CDA. Yan et al.^5^ developed a method (DWNN-RLS) to predict circRNA-disease associations based on regularized least squares of Kronecker product kernel. These models have the disadvantage of requiring high-confidence negative samples, and may need to retrain the models for new circRNAs and diseases. Second, network-based methods mainly utilize the network structure information between circRNAs and diseases, such as circRNA functional similarity, disease semantic similarity, circRNA-target gene interaction, etc., to predict potential circRNA-disease associations. For example, KATZHCDA^6^ adopted the KATZ algorithm to infer CDA. Huseyin et al.^7^ used random walk with restart (RWR) to predict CDA. Zhang et al.^8^ identified potential CDA by constructing similarity networks and using the linear neighborhood label propagation method (CD-LNLP). Wang et al.^9^ proposed a computational method (PreCDA) based on a graph-based recommendation algorithm to predict CDA. Lei et al.^10^ proposed a computational model based on similarity network fusion (GBDTCDA) to predict CDA. However, these methods have the drawback of requiring high-quality biological network models, and may lack sufficient network information for new circRNAs and diseases.Third, deep learning methods use deep neural networks to learn the complex nonlinear relationship between circRNAs and diseases, and can integrate multiple heterogeneous data sources. These models have the advantage of automatically extracting high-level features from data, and capturing deep association patterns. For example, Deepthi et al.^11^ used deep autoencoder (AERF) to identify CDA. Niu et al.^12^ used graph Markov neural network (GMNN2CD) to capture CDA information and make predictions. Lu et al.^13^ used a multi-layer neural network for deep matrix factorization method (DMFCDA) to predict unknown CDA. Lan et al.^14^ designed a knowledge graph with attention mechanism (KGANCDA) to infer unknown CDA. Wu et al.^15^ designed a transformer-based encoder (KGETCDA) to infer CDA. A disadvantage of them is that they mostly require embedding large amounts of labeled data to train the network parameters, cannot capture deep information through explicit methods, and may need additional data augmentation or transfer learning techniques to deal with new circRNAs and diseases.

While current models have yielded promising results, there are still some limitations. Given the scarcity of data concerning the association between circRNA and diseases (CDA), existing models rely on reasoning directly over triplets comprising embedded entities and relations^16^, without the explicit capture of local information^17^. This poses a challenge for accurate CDA prediction. Consequently, the proposition of a method that harnesses multi-source data and explicitly extracts local depth information from triplets holds considerable significance.

In this paper, we introduce a novel CDA prediction model: KGRACDA. Our model is based on a knowledge graph with recursive and attention aggregation, which can effectively utilize multi-source heterogeneous datasets and explicitly capture the local deep details between circRNA and disease. Firstly, we use a multi-source heterogeneous ncRNA dataset, which includes circRNA, miRNA, lncRNA and disease entities, as well as their associations. Then, we construct a directed knowledge graph of circRNA, lncRNA, miRNA and disease associations. Finally, we use a recursive method to build a graph neural network, and use attention to aggregate the local deep information of the knowledge graph, to obtain reliable CDA matching scores. Experiments show that KGRACDA outperforms other state-of-the-art (SOTA) models. Moreover, we also develop a web platform, where users can easily use KGRACDA to predict potential CDAs.

## RESULTS

### Overview of KGRACDA

KGRACDA is a model for predicting circRNA-disease association (CDA), which supports end-to-end CDA analysis. Unlike existing methods that mainly use feature vector embeddings, it utilizes knowledge graph techniques to represent circRNA, miRNA, lnRNA and diseases as entities and relations. This representation can explicitly capture and mine the local and deep information in the knowledge graph, thus better predicting the relationships between circRNA and diseases. Importantly, KGRACDA uses a recursive method to build a graph neural network, which enables the model to capture the local information between nodes from shallow to deep, fully aggregate the relations between entities, select strongly associated nodes with attention mechanism, and obtain reliable CDA prediction scores.

### Datasets

In this study, we utilize a total of three datasets. Firstly, we have gathered more than 300 circRNAs, 265 miRNAs, and 400 lncRNAs from previous research^14^, comprising over 1000 distinct entities, including 79 disease types. Concurrently, we establish the relationships between these entities, specifically encompassing over 340 circRNA-disease, more than 100 miRNA-disease, approximately 500 lncRNA-disease, around 140 circRNA-miRNA, and nearly 200 miRNA-lncRNA interactions, summing up to 1377 relationships. These entities and relationships collectively form Dataset 1.

The second dataset is a multisource dataset that includes a diverse array of RNA and disease entities and their corresponding relationships. It consists of 561 circRNAs, nearly 660 miRNAs, over 1000 lncRNAs, and a total of 190 disease types, resulting in almost 2500 distinct entities^15^. This dataset is remarkable for its extensive network of relationships, exceeding 25,500 interactions. Specifically, it encompasses approximately 1400 circRNA-disease, over 10,000 miRNA-disease, nearly 3300 lncRNA-disease, more than 1100 circRNA-miRNA, and over 9500 miRNA-lncRNA relationships. Experimenting with such a large dataset significantly enhances the accuracy and reliability of our experimental results.

Additionally, we incorporated Dataset 3 from prior research^14^. Dataset 3 consists of 514 circRNAs, over 660 miRNAs, more than 540 lncRNAs, and a total of 62 disease types, culminating in a grand total of over 1800 distinct entities. We concurrently establish the relationships between these entities, which includes nearly 650 circRNA-disease, over 730 miRNA-disease, more than 1000 lncRNA-disease, approximately 750 circRNA-miRNA, and over 300 miRNA-lncRNA interactions, totaling 3059 relationships. These entities and their associated relationships constitute Dataset 3.

To provide a visual overview of the dataset’s composition and distribution, we have depicted Figure 1, which illustrates the associations between circRNAs and diseases in the dataset. Figure 1A details the number of entities in each of the three datasets. As evident from the Figure 1B, the connections between circRNA and diseases are relatively sparse, with most entities having a small degree. From a knowledge graph perspective, this represents a low-density graph with few relationships between nodes and considerable blank and empty space. We use a circular bar chart to visualize the relationship between several specific circRNAs in the data set and disease, as shown in Figure 1C. We further analyze the datasets and visualized the distribution of relationships, degree statistics of diseases and circRNAs in the three datasets. For details, please refer to Supplementary Figures S1-S3.

**Figure 1.**
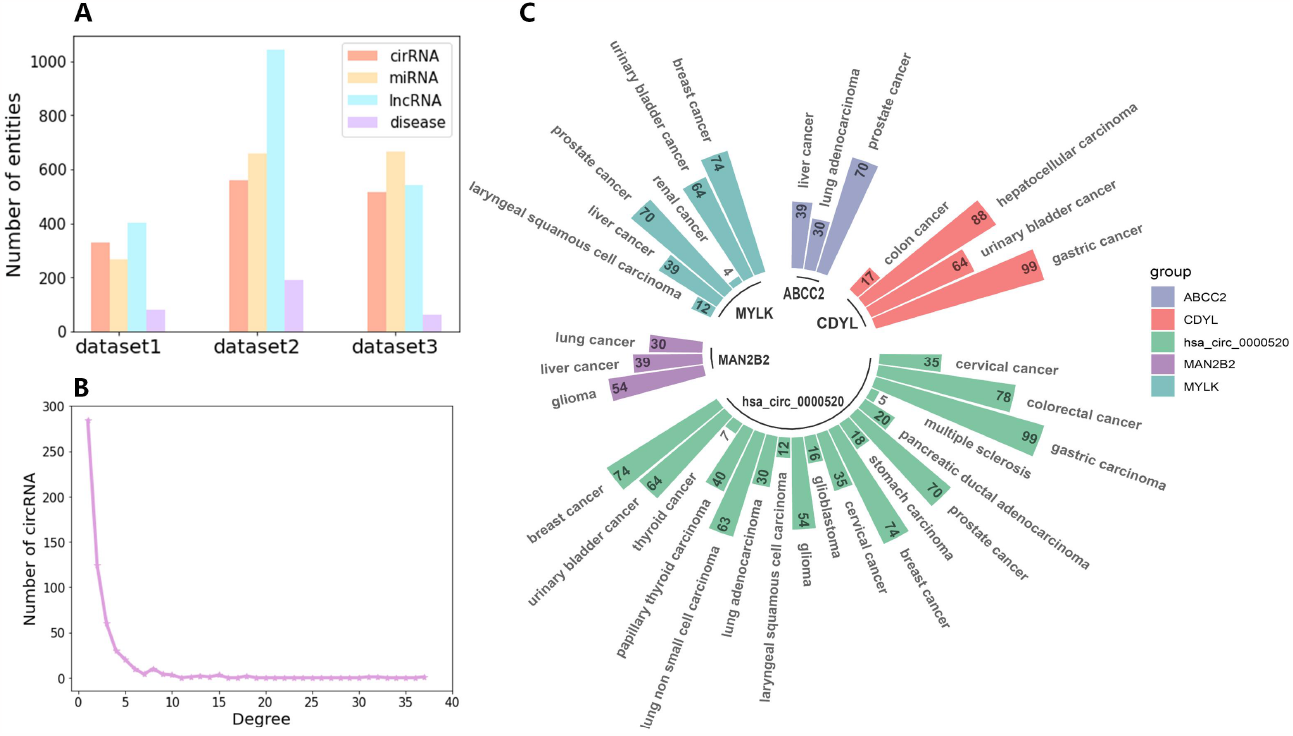
The visualization of datasets. **(A)** Distribution of the number of entities in three datasets. **(B)** The degree distribution of dataset2. It represents the number of corresponding circRNA associating with different numbers of disease in dataset2. **(C)** Relationship between some circRNAs and diseases in dataset 2.

We conduct a series of experiments on three datasets— dataset1, dataset2, and dataset3—to demonstrate the higher accuracy of KGRACDA in predicting CDA. First, we introduce the evaluation metrics used in our experiments. To enhance persuasiveness, we compare the KGRACDA method with other state-of-the-art approaches (SOTA) based on these evaluation metrics. Additionally, we perform ablation experiments to further assess the effectiveness of the KGRACDA model and identify its optimal parameter settings. Finally, we evaluate the ability of KGRACDA to identify potential associations between circular RNAs and diseases by providing specific examples of disease-circRNA prediction relationships. Through these experiments, we demonstrate that KGRACDA, by explicitly capturing local deep-level details, achieves higher accuracy in predicting CDAs.

### Evaluation criteria

We evaluate KGRACDA using 5-fold cross-validation and compare it with other state-of-the-art models. The 5-fold cross-validation method fully utilizes all samples in training, which helps reduce overfitting and make the model training more effective. In 5-fold cross-validation, we randomly divide all samples of the entire dataset into five parts, and select each part of the dataset as the test dataset, while the remaining four parts of the dataset as the training dataset. Then, by testing on each fold of data, we obtain the prediction scores of diseases and circRNA, and sort the prediction scores in descending order. Finally, by averaging the results of each fold, we obtain the average results of the five folds. Finally, we use some typical evaluation metrics to evaluate the model results, including precision, accuracy, recall, and F1 score.

The mathematical expressions of these evaluation metrics are as follows:

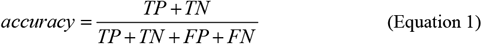

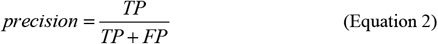

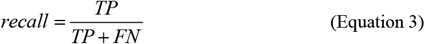

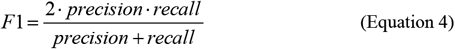

Where TP represents the number of positive samples correctly predicted, TN represents the number of negative samples correctly predicted; FP represents the number of positive samples incorrectly predicted, FN represents the number of negative samples incorrectly predicted. We use the area under the ROC curve (AUC) and the area under the PR curve (AUPR) to evaluate the model. The PR curve is a curve drawn with recall as the horizontal axis and precision as the vertical axis. AUPR is an effective indicator that objectively reflects the performance changes of the model, reflecting the model’s ability to identify positive samples at different thresholds. The ROC curve is a curve drawn with the false positive rate (FPR) as the horizontal axis and the true positive rate (TPR) as the vertical axis. AUC can be used to represent the specificity and sensitivity of the model, reflecting the model’s classification effect on positive and negative samples at different thresholds. Where FPR represents the proportion of samples that are actually negative but are incorrectly judged as positive among all samples that are actually negative. TPR represents the proportion of samples that are actually positive and are correctly judged as positive among all samples that are actually positive. Their mathematical expressions are as follows:

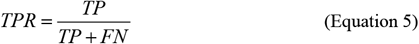

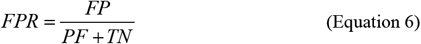

In addition, we further measure the predictive power of the model using the number of correctly identified CDA, expressed as *Top* − *k* (*k* = 10, 20,30, 40).

To demonstrate the superiority of the KGRACDA method, we compare KGRACDA with nine current best models, including network-based (CD-LNLP^8^, KATZHCDA^6^ and RWR^7^), traditional machine learning-based (DMFCDA^13^ and RNMFLP^4^) and deep learning-based (AE-RF^11^, GMNN2CD^12^, KGANCDA^14^ and KGETCDA^15^) methods. Below, we systematically compare KGRACDA with the above nine current best methods from two aspects: experimental evaluation metrics and hypothesis testing. Under the same experimental environment, using the same experimental data, we obtain the final results of the above 10 models on 3 datasets with 5-fold CV. The parameters of the other nine methods are consistent with the original papers.

For the relatively small datasets dataset1 and dataset3, as depicted in Figures 2A-2B and 4A-4B, KGRACDA outperforms other models in terms of precision, accuracy, AUC and AUPR, despite the sparse nature of the data. These results underscore our model’s excellent predictive capability for CDAs. Detailed information can be found in Supplementary Table S1. On dataset1, KGRACDA achieves an accuracy of 0.4988, a recall rate of 0.9291, an AUC of 0.9312, and an AUPR of 0.0359. Similarly, on dataset3, KGRACDA achieves an accuracy of 0.4978, a recall rate of 0.8450, an AUC of 0.8602, and an AUPR of 0.0609. Furthermore, to further evaluate the model’s predictive ability, we compute the top-10, top-20, top-30, and top-40 correct predictions for each of the ten models mentioned above (as shown in Figures 2C and 2C). Experimental results demonstrate that KGRACDA exhibits superior overall predictive performance compared to other models, once again highlighting its exceptional capacity to capture local deep-level information encoded in knowledge graph triplets. Additional details are provided in Supplementary Table S2.

**Figure 2.**
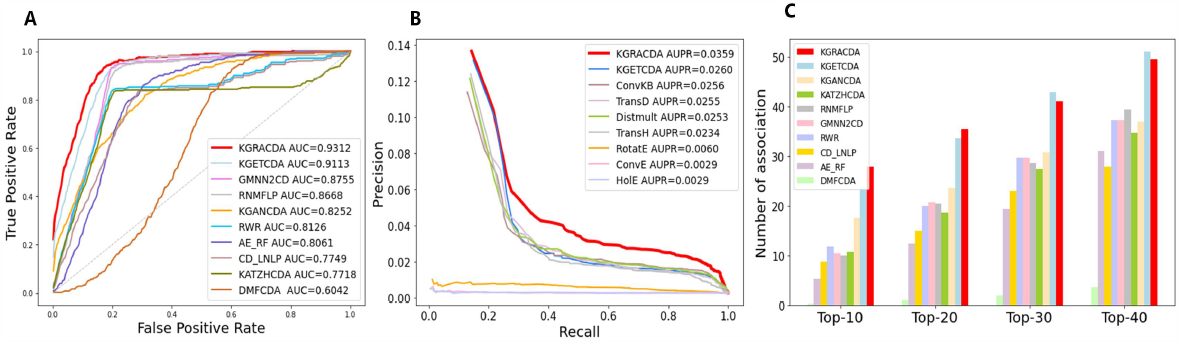
The comparison of KGRACDA with other predicting methods on dataset1. **(A)** The AUC comparison (**B**) The AUPR comparison (**C**) The comparison of the number of correctly identified CDA

For the larger heterogeneous dataset dataset2, as illustrated in Figures 3A and 3B, KGRACDA continues to outperform the other nine models in terms of precision, F1 score, accuracy, AUC and AUPR. This reaffirms KGRACDA’s explicit ability to capture local triplet details—a capability not shared by other models. Similarly, we compute the top-k metrics for the ten models on dataset2, as shown in Figure 3C. The comprehensive performance of KGRACDA on dataset2 remains superior to other state-of-the-art (SOTA) methods. Upon further analysis of the experimental results, we attribute this performance to the complexity and heterogeneity of data in dataset2, which makes relationship discovery more intricate. While other SOTA methods face limitations due to these complexities, KGRACDA still exhibits strong predictive capabilities. This resilience stems from its capacity to mine and capture local depth details within knowledge graph nodes.

**Figure 3.**
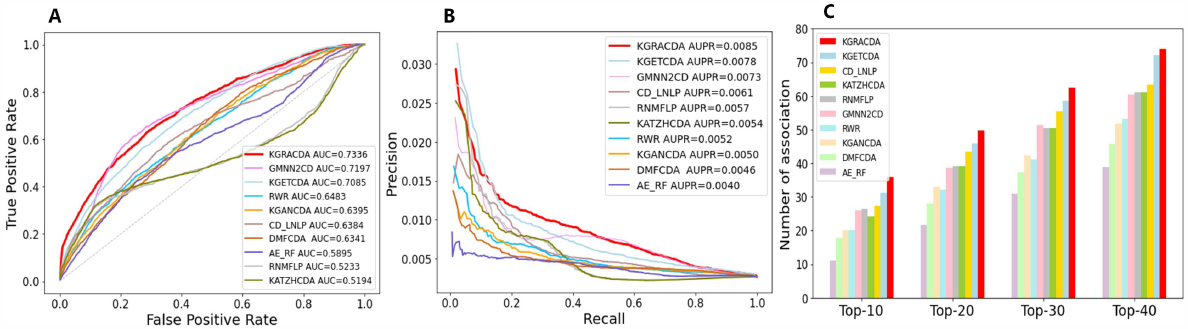
The comparison of KGRACDA with other predicting methods on dataset2. **(A)** The AUC comparison (**B**) The AUPR comparison (**C**) The comparison of the number of correctly identified CDA

**Figure 4.**
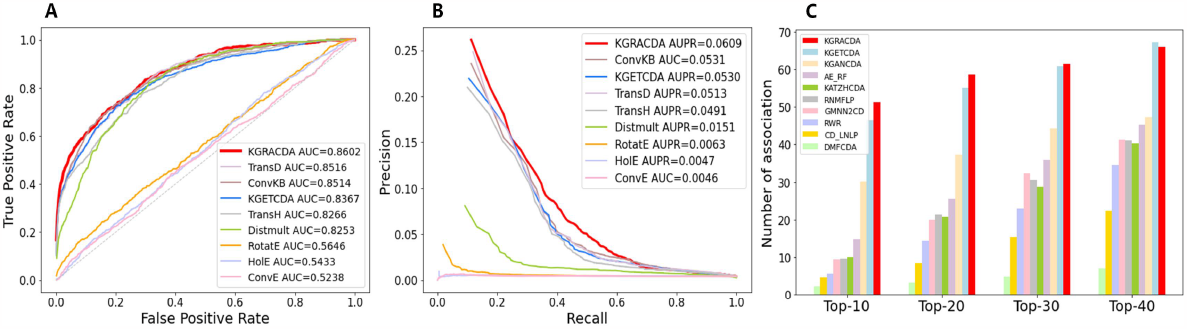
The comparison of KGRACDA with other predicting methods on dataset3. **(A)** The AUC comparison (**B**) The AUPR comparison (**C**) The comparison of the number of correctly identified CDA

### Statistical hypothesis test

To further evaluate the significant difference between KGRACDA and other SOTA methods from a statistical perspective, we adopt the non-parametric Wilcoxon signed-rank test method^18^. The non-parametric Wilcoxon signed-rank test method is a statistical method for comparing whether the medians of two related samples have significant differences, which does not require the assumption that the data follows a normal distribution or other specific distributions, and at the same time, it is not affected by outliers or extreme values. This method can effectively utilize the size of the difference, thereby effectively detecting the statistical difference between the two models. We set the null hypothesis (*H*_0_) as no significant difference between KGRACDA and the baseline (*H*_0_ :difference=0), the alternative hypothesis (*H*_1_) as *H*_1_ : difference ≠ 0, and set the default p-value threshold to 0.05. As shown in Supplementary Table S3, the p-values between KGRACDA and other models are all less than 0.05, so we reject the null hypothesis. Therefore, from a statistical point of view, KGRACDA is superior to other SOTA methods.

### Ablation Study

We conduct ablation experiments to further analyze the KGRACDA model. We first compare the performance of KGRACDA with other knowledge graph models. Then, we discuss the impact of different graph neural network layers, attention layer dimensions on the results. Finally, we study the effect of learning rate, learning rate decay coefficient and weight decay coefficient on the performance of KGRACDA model. Here we show the results on dataset1(shown in Figure 5), while the results on dataset2 and dataset3 are available in the Supplementary Figures S4-S5.

**Figure 5.**
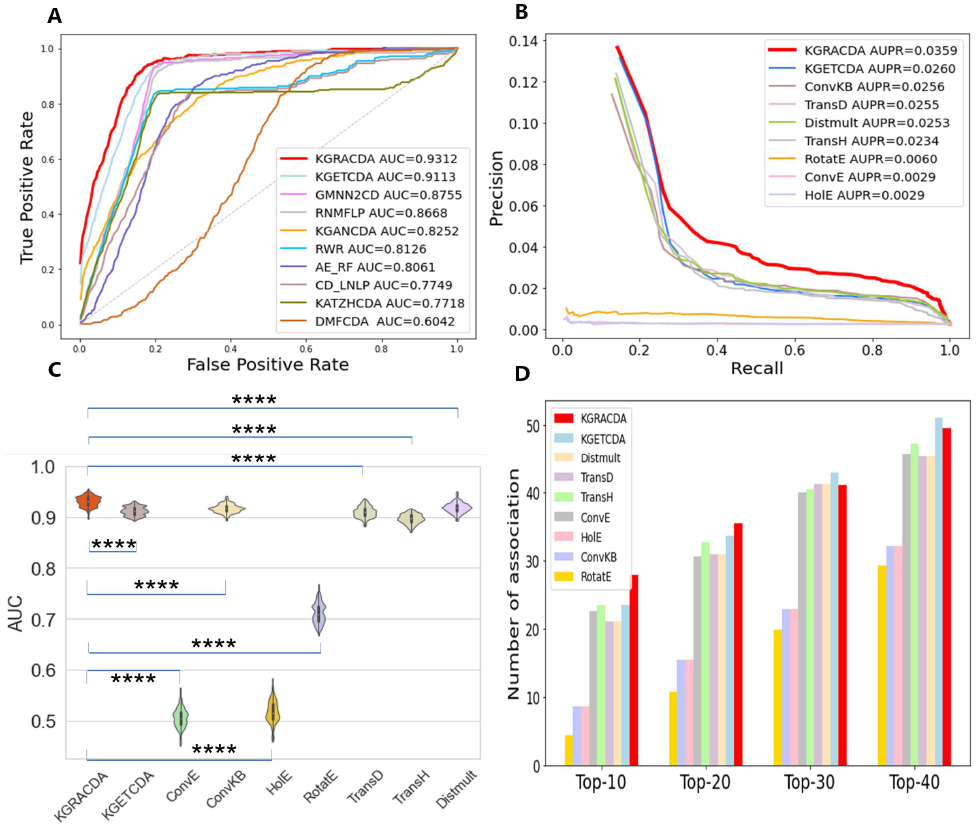
The comparison of KGRACDA with other KG methods on dataset1. **(A)** The AUC comparison (**B**) The AUPR comparison **(C)** The AUC indicators of multiple experiments are visualized in the form of violin graphs, marked with the level of significance difference for statistical hypothesis testing (**** for P-values<0.0001). **(D)** The comparison of the number of correctly identified CDA

To better validate the effectiveness of the KGRACDA model, we compare its performance with other knowledge graph models. In this study, we compare KGRACDA against a total of eight KG models: TransD^19^, TransH^20^, ConvKB^21^, ConvE^22^, KGETCDA^15^, HolE^23^, RotatE^24^, and Distmult^25^. We evaluated their performance in predicting CDA on dataset1 using a 5-fold cross-validation approach, and the results for the area under the curve (AUC) are shown in Figure 5A and 5C. The AUC obtained by the KGRACDA model is slightly higher than that of KGETCDA, Distmult, TransD, ConvKB, and TransH, and significantly higher than that of ConvE, HolE, and RotatE, demonstrating the effectiveness of our model. The comparison results of AUPR and Top-k indexes are similar to those of AUC. For details, please see Figure 5B and 5D. Specific data are presented in Supplementary Table S4-S5, and the results of datasets2 and dataset3 are shown in Supplementary Figure S4-S5.

Similarly, we use the non-parametric Wilcoxon signed-rank test to evaluate the statistically significant differences between KGRACDA and other KG methods. As shown in Supplementary Table S6, the P-values of KGRACDA and other KG models were all less than 0.05, so we rejected the null hypothesis. Therefore, from a statistical point of view, KGRACDA is superior to other KG methods.

We further investigate the impact of graph neural network (GNN) layers and attention layer dimensions on the results. Here we present the results of dataset1. Figure 6A shows the changes in AUC,AUPR, and time as the number of GNN layers increases. Figure 6B shows the change of model AUC under the two variables of GNN layer number and attention layer dimension. From Figure 6, it is evident that when the GNN layers are set to 32 and the attention layer dimensions are also set to 32, the KGRACDA model achieves optimal performance. At this configuration, both AUC and AUPR evaluation metrics exhibit high values, and notably, the training time is shorter compared to using more GNN layers or larger attention layer dimensions.

**Figure 6.**
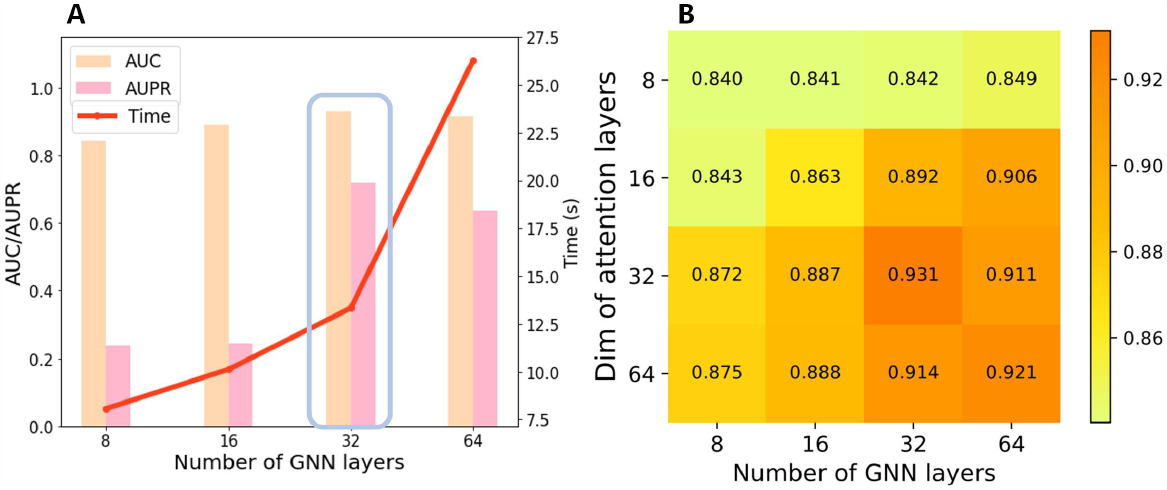
The results of ablation experiments on dataset1. **(A)** The effect of different GNN layers dimensionality. We select 32 here after considering both AUC and AUPR performance, and time consumption. For better visualization, AUPR is uniformly stretched to better highlight the trend of increase and decrease. (**B**) The effect of the number of GNN layers and the dim of attention layers.

Increasing the number of GNN layers from 8 to 32 results in improved model performance, indicating that within a certain range, more GNN layers effectively capture information from relationships. However, when the number of GNN layers increases from 32 to 64, KGRACDA’s performance no longer improves and may even decline. Simultaneously, training time significantly increases. This phenomenon could be attributed to excessive GNN layers leading to sparse feature representations, potentially introducing irrelevant information or noise^26^. Similar observations apply to evaluating attention layer dimensions. For the results of dataset2 and dataset3, please refer to Supplementary Figure S6-S7.

### Model interpretability

This paper provides two methods to explain how KGRACDA captures the local depth information of triplets from the knowledge graph and makes predictions. First, we provide a visual method to demonstrate KGRACDA’s ability to acquire local information. We obtain 300 known circRNA-disease relationships and 300 unknown circRNA-disease relationships from dataset1. We consider the known relationships as positive pairs and the unknown relationships as negative pairs, and apply the PCA method^27^ to visualize the high-dimensional features of entities in the knowledge graph in two dimensions. As can be seen from Supplementary Figure S8A the distribution of positive pairs and negative pairs is very random and irregular in the initial state. After KGRACDA’s processing, the spatial positions of positive pairs and negative pairs are basically separated, and there is a clear clustering phenomenon in the distribution. The positive pairs with large initial differences verified by biological experiments are clustered together after KGRACDA, which clearly shows how KGRACDA captures the depth information in the knowledge graph and extracts features from entities (see Supplementary Figure S8B). We focus on the aggregation results of three specific categories of diseases: systemic lupus erythematosus (SLE), hepatocellular carcinoma, and lung cancer. As can be seen from Supplementary Figure S8B, the KGRACDA model aggregated positive pairs of these three diseases together, achieving the role of distinguishing positive pairs from negative pairs. Second, we randomly select 10 circRNAs and 10 diseases from the dataset, and use the KGRACDA model to predict the association score between each circRNA and each disease. We visualize this association score matrix to the heatmap in Supplementary Figure S9, which can intuitively find that after KGRACDA model’ s feature extraction, the relationship between circRNA and disease becomes clear and straightforward. KGRACDA rarely has ambiguous situations for CDA prediction, reflecting the model’s powerful prediction ability. Taking squamous cell carcinoma as an example, after KGRACDA’s prediction score, cMras and circ-ITCH have significantly higher scores than other circRNAs, and the fact that these two circRNAs are related to squamous cell carcinoma has been supported by literature^28,29^, indicating that KGRACDA can fully utilize the local structure of the “squamous cell carcinoma” node, and has accuracy in capturing information. Through the above analysis, we can prove the interpretability of KGRACDA model.

### Parameter setting and sensitivity analysis

We implement KGRACDA in PyTorch using NVIDIA Titan Xp GPU, and optimize the whole model using Adam optimizer^30^ and Exponential LR learning rate scheduler^31^. The learning rate was set to 2e-4, the learning rate decay rate was set to 0.999, the regularization decay weight was set to 1e-3, and the number of training epochs was 50. In order to make full use of the data, we set the attention layer dimension to 32, and set the graph neural network to 32 layers. To reduce the impact of overfitting on the model, we set dropout to 0.02. In addition, the detailed information of parameter settings is shown in the Supplementary Table S7.

We conduct a parameter sensitivity study on the learning rate, the learning rate decay coefficient, and the regularization weight decay coefficient, and reported the impact of these parameters on the model performance (AUC) in Supplementary Table S8. The experimental results show that when the parameters vary within a certain range, the overall value of AUC presents similar results, reflecting the robustness of KGRACDA.

### Case studies and survival analysis

To further evaluate KGRACDA’s efficacy, we conduct individual case studies on the three distinct datasets. We train the PKGRACDA model using known CDA, and then predict the potential probabilities for all unknown CDAs and rank them in a descending order based on these probabilities. In addition, we meticulously gather experimental evidence and validate these predictions by cross-referencing them with information available in publicly accessible databases and recent literature. The results are presented in the accompanying table. For dataset1, we select systemic lupus erythematosus (SLE), a disease that previous research has demonstrated to benefit from circRNA investigations^32^. From Table 1, it is evident that among the top 10circRNAs related to SLE predicted by KGRACDA, seven of them have been confirmed by existing studies, namely circptpn22, hsa_circ_0000479, hsa_circ_0045272, hsa_circ_0057762, hsa_circ_0049224, has_circ_0049220, hsa_circ_0008945. For example, hsa_circ_0000479 serves as a novel biomarker for SLE^33^, and hsa_circ_0049224 is downregulated in SLE patients^34^.

**Table 1.**
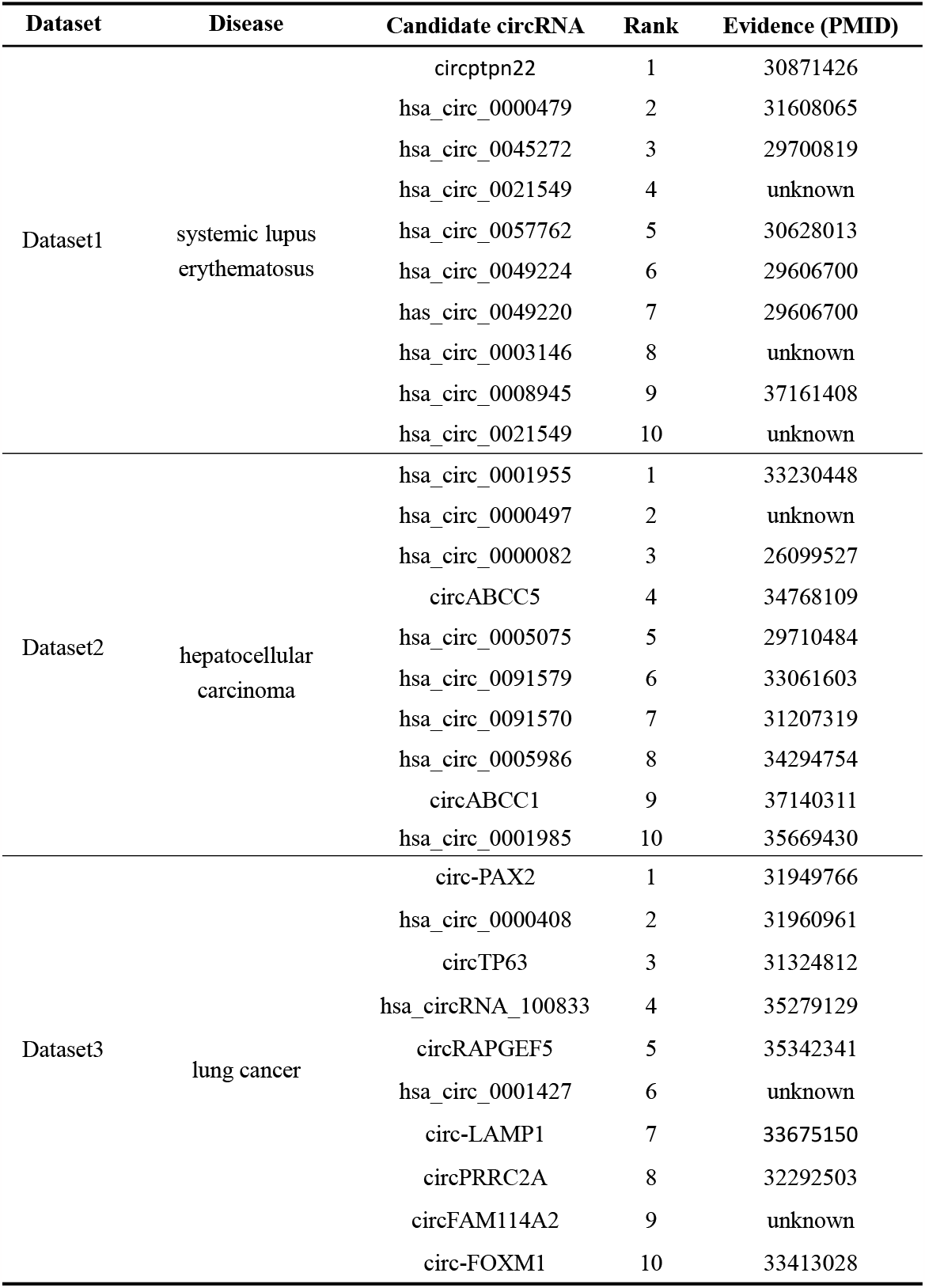
The top-10 predicted systemic lupus erythematosus, hepatocellular carcinoma, or lung cancer related candidate circRNAs.

For dataset2, we choose hepatocellular carcinoma, a grave ailment with substantial impact on human health. Utilizing the KGRACDA model, we identify the top 10 liver cancer-related circRNAs based on the scores as shown in Table 1. Remarkably, seven of these circRNAs were corroborated by existing literature: hsa_circ_0001955, hsa_circ_0000082, hsa_circ_0005075, hsa_circ_0091579, hsa_circ_0001570, hsa_circ_0005986, hsa_circ_0001985. For example, hsa_circ_0005075 exhibited significant expression disparities in hepatocellular carcinoma (HCC) tissues^35^, while hsa_circ_0091579 is identified as a diagnostic and prognostic marker for HCC^36^.

For dataset3, we chose lung cancer as the predicted disease. The KGRACDA model predicted the top 10 lung cancer-related circRNAs based on the scores as shown in Table 1, among which eight circRNAs found validation in existing literature, namely circ-PAX2, hsa_circ_0000408, circTP63, hsa_circRNA_100833, circRAPGEF5, circ-LAMP1, circPRRC2A, circ-FOXM1. Notably, circ-PAX2 is found to promote lung cancer cell proliferation and metastasis by sponging miR-186^37^, and circ-FOXM1 is reported to inhibite the development of non-small cell lung cancer by regulating the miR-149-5p/ATG5 axis^38^. We have visualized the model’s predictions in a circular bar graph, which can be found in Supplementary Figure S10.

These findings underscore the predictive precision of the KGRACDA model and its potential in advancing our understanding of circRNAs in the context of SLE, liver cancer, and lung cancer biology. Furthermore, the CDAs predicted by KGRACDA but lacking current literature support hold promise for further scrutiny through biological experiments. These unverified predictions represent fertile ground for expanding our understanding of circRNAs in various disease contexts.

In addition, to comprehensively evaluate the prediction performance of the KGRACDA model, we leverage publicly available TCGA liver cancer data to perform survival analysis on the top 10 liver cancer-related circRNAs predicted by KGRACDA. The results of survival analysis are shown in Supplementary Figure S11. We find that the expression levels of the circRNAs predicted by KGRACDA were highly correlated with disease progression. This dual validation approach reinforces the utility of KGRACDA in not only identifying CDAs but also shedding light on their clinical relevance and prognostic significance, thus emphasizing its significance in the realm of circRNA research.

In light of the lack of circRNA transcriptome sequencing data, we employ an alternative approach by utilizing the host genes of the predicted circRNAs for conducting survival analysis. This analysis is carried out using data from TCGA in conjunction with the GEPIA database(http://gepia2.cancer-pku.cn)^39^. Based on the expression differences of the host genes of the circRNAs predicted by KGRACDA in dataset2, we generate the survival analysis curves of the host genes (ABCC1, AXIN1, CDKN2A, ABCC5) corresponding to four selected circRNAs (i.e., circABCC1, hsa_circ_0001985, hsa_circ_0000082, circABCC5) as shown in Supplementary Figure S11. Taking Figure S11D as an example, patients are divided into high-expression and low-expression groups according to the expression level of ABCC5 gene. Subsequently, we perform survival analysis using Kaplan-Meier (KM) and Log-rank-type test ^40^. The results reveal a significant statistical difference in survival time between the two groups (95% CI and log-rank P value < 0.05), indicating that ABCC5 expression is significantly associated with liver cancer prognosis. The survival analysis conducts for the other three genes yielded similar results.

In summary, following rigorous experimental validation, it has become evident that the expression levels of the circRNAs predicted by KGRACDA are highly correlated with the progression of the disease, indicating their potential as prognostic markers. These findings open up new possibilities for subsequent medical treatment.

## WEB-BASED VISUALIZATION AND PREDICTION

We have updated an online interactive CDA prediction platform HNRBase v2.0, as shown in Figure 7. HNRBase v2.0 builds on HNRBase, adding rich circRNA data visualization.

**Figure 7.**
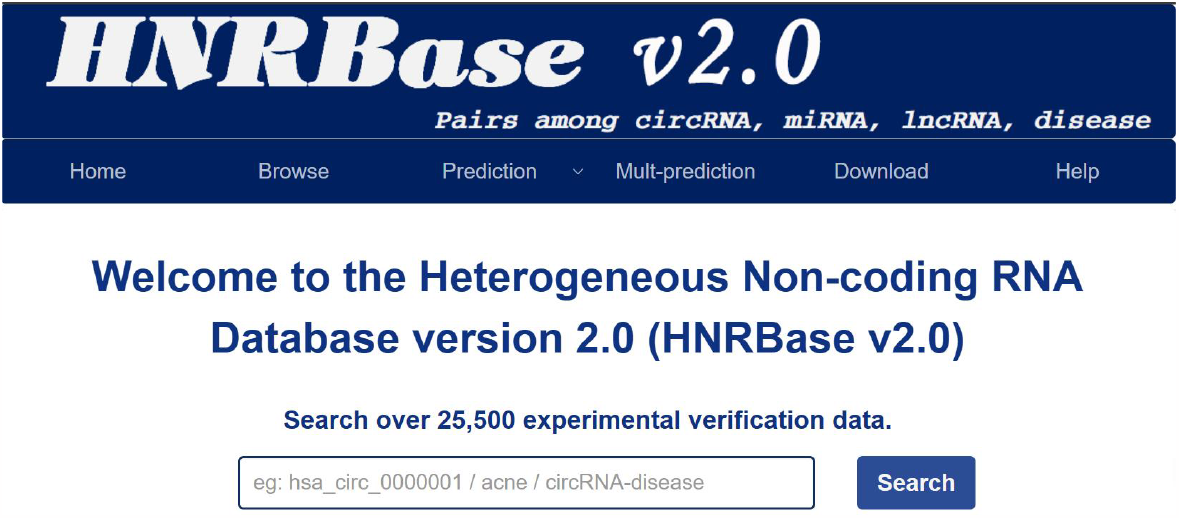
The homepage of HNRBase v2.0 platform.

First, HNRBase v2.0 adds a many-to-many prediction function, that is, users can input multiple circRNAs and multiple diseases at one time, and the website will display a heat map of the CDA prediction scores predicted by the KGRACDA model. This visualization method is fast and convenient for users to obtain CDA prediction scores online. An example of many-to-many CDA prediction is shown in Supplementary Figure S12A. Secend, HNRBase v2.0 also adds the visualization of circRNA data sets, as shown in Supplementary Figure S12B. Through the dynamic pie chart and bar chart, the composition of the data set is clearly shown. Moreover, HNRBase v2.0 adds information related to circRNA, which facilitates users to query circRNA data more deeply. HNRBase v2.0 adds the igv web embedding tool, which allows users to query the genomic information of circRNA without leaving the web page by clicking “useigv”. An example of using the igv web embedding tool^41^ is shown in Supplementary Figure S13A. In addition, HNRBase v2.0 sets the word cloud as a link, and users can jump to Google search by clicking on a specific entity in the word cloud, which facilitates users to conduct deeper search on the entity. A specific example is shown in Supplementary Figure S13B.

In summary, HNRBase v2.0 provides more functions and convenience for users, allowing them to obtain, process, analyze and use data more efficiently, and facilitate further research in the field of CDA prediction.

## DISCUSSION AND CONCLUSION

Research has shown a strong correlation between circRNAs and human diseases. Exploring the associations between circRNAs and diseases can offer valuable insights and methods for understanding disease mechanisms, facilitating diagnosis, and devising effective treatment. Such research bears substantial significance in the realms of disease prevention and management.

However, due to the limited availability of circRNA-disease association data, existing computational methods still have room for improvement in capturing local deep-level features related to CDAs. In this paper, we introduce the KGRACDA model—a novel approach based on graph neural networks for CDA prediction within knowledge graphs. KGRACDA is designed to explicitly capture local triplet information and effectively predicts potential CDAs. We evaluated the performance of the KGRACDA model using three distinct datasets, and our experimental results demonstrate its superiority over state-of-the-art (SOTA) methods. Notably, it excels in extracts intricate local deep-level features from biological knowledge graphs.

Additionally, we have developed an interactive online platform for CDA prediction. This platform provides users with friendly interfaces for visualizing data, conducting searches, download information, and accessing model predictions concerning CDAs. By presenting results in this manner, we empower users with enhanced functionality and convenience, enabling efficient data acquisition, processing, analysis, and utilization. Furthermore, this platform contributes to further research in the field of CDA prediction.

However, our model still has some limitations. Our model has limited ability to utilize multimodal data. Although we create a heterogeneous knowledge graph using multiple sources of data sets, we have limited assistance in utilizing biological knowledge. If we can integrate more biological knowledge into the model, it is likely to improve the model’s ability to mine local information. Therefore, we will try to introduce RNA sequences and gene expression information into the model in our future research, to help improve the model’s performance in predicting CDA.

## STAR☆METHODS

In this section, we present the KGRACDA method, designed to effectively capture localized deep information. Figure 8 provides an overview of the KGRACDA model, which comprises three key components: knowledge graph construction, recursive creation of graph neural networks, and attention aggregation for deriving prediction scores.

**Figure 8.**
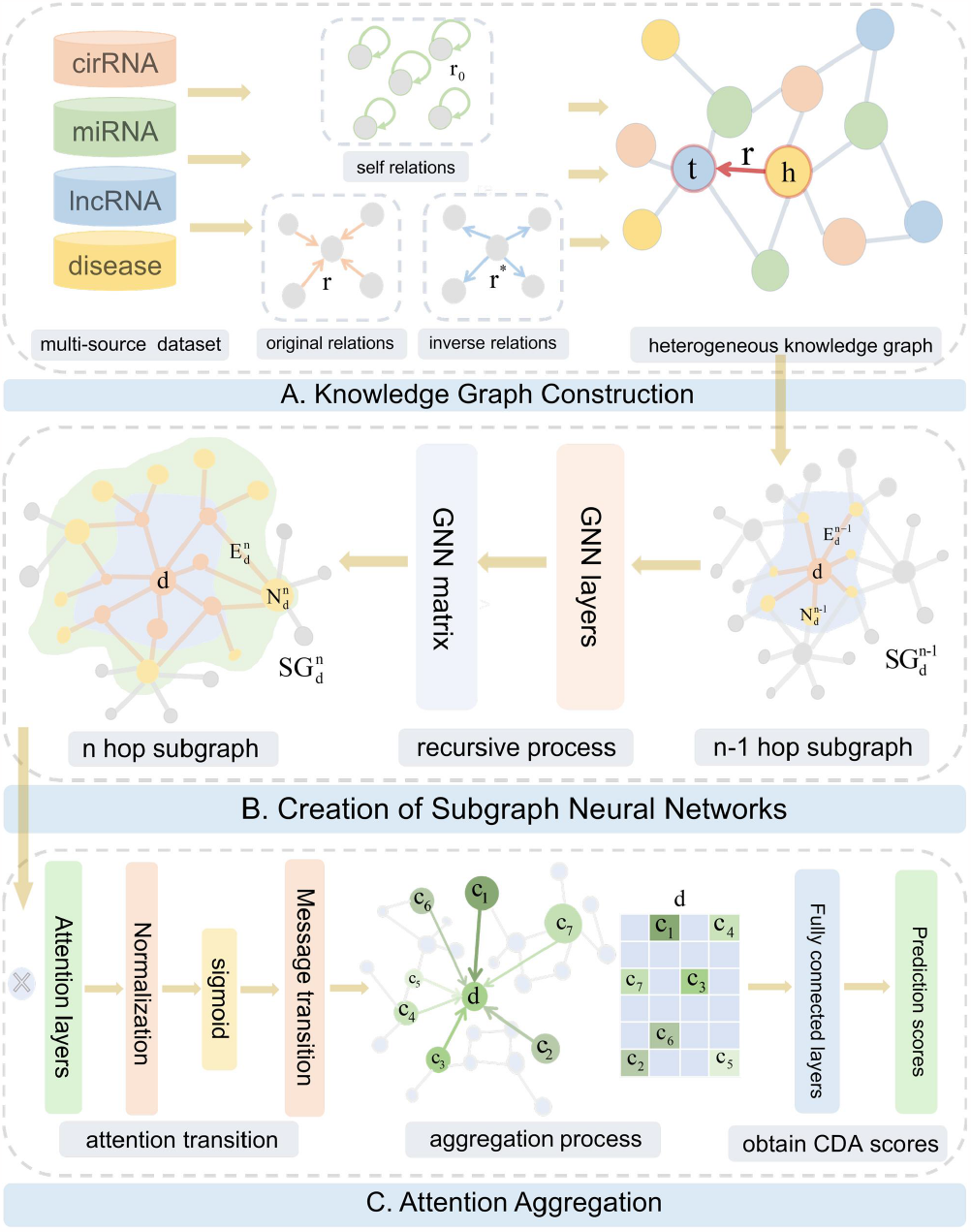
The flowchart of KGETCDA. **(A)** Construct the knowledge graph with directed self-loop. **(B)** Subgraphs are constructed by GNN. **(C)** CDA prediction scores are obtained by attention aggregation.

### A. Knowledge Graph Construction

In recent years, the knowledge graph, initially introduced by Internet companies to enhance search engine functionality, has seen increasing application for handling complex, multi-source, and heterogeneous data^42^. A Knowledge graph offers a structured representation of concepts, entities, and their interrelationships within the real world, presenting information in a structure that closely aligns with human cognitive processes., It offers a more effective framework for organization, management, and comprehension of vast amounts of data^43^. Moreover, knowledge graph serves the purpose of modeling, identifying, discovering and inferring intricate connections between entities and concepts. It functions as a computable model for understanding the interconnections among various elements and has seen wide application in fields such as search engines, language comprehension, visual scene analysis, and beyond^44^. What sets the knowledge graph apart is its ability to facilitate data integration, supporting the generation of new knowledge and establishing connections between data points that might have remained unrealized before^45^. Encouraged by these prospects, we have chosen to embrace the knowledge graph methodology to address the challenge of predicting the associations between circRNA and diseases.

In this investigation, we have developed a knowledge graph (KG) focusing on circRNA, lncRNA, miRNA and diseases, utilizing the dataset introduced in the preceding section. The KG is formally denoted as *KG* = (*E,V, R*), where *V* represents the set of nodes in the *KG*, with each element representing a circRNA, miRNA, lncRNA or disease under consideration. The set *R* signifies the relations within the *KG*, specifically circRNA-disease (*circ* − *d*), miRNA-disease (*mi* − *d*), lncRNA-disease (*lnc* − *d*), circRNA-miRNA (*circ* − *mi*), and miRNA-lncRNA (*mi* − *lnc*), denoted as *R* = {*circ* − *d, mi* − *d, lnc* − *d, circ* − *mi, mi* − *lnc*}. *E*, the set of edges in the *KG*, is composed of triplets in the form *E* ={(*h, r, t*) | *h, t* ∈*V, r* ∈ *R*}, where *h* signifies the head entity, *r* represents the relation, and *t* is the tail entity. Through the construction of the *KG*, the research problem is reformulated to derive the prediction score of (*d, d* − *circ, circ*), where *circ* and *d* denote any circRNA and disease within the dataset, respectively. Subsequently, we rank these prediction scores in descending order to anticipate the most pertinent circRNAs associated with a given disease.

During the knowledge graph construction, depicted in Figure 8A, we utilize not only the triplets corresponding to the original relations but also those associated with the inverse relations and self-relations. Given a triplet (*e*_1_, *r, e*_2_), its inverse relation triplet is represented as (*e*_2_, *r*^*^, *e*_1_), and its self-relation triplet includes (*e*_1_, *r*_0_, *e*_1_) and (*e*_2_, *r*_0_, *e*_2_), where *r*^*^ denotes the inverse relation of *r*, and *r*_0_ signifies the self-relation. This approach to knowledge graph construction, to a certain extent, mitigates the issue of data sparsity in the CDA dataset and facilitates the model in extracting local deep information from the knowledge graph.

### B. Recursive Creation of Graph Neural Networks

The Graph Neural Network (GNN) is considered an effective approach for solving node-level, edge-level, and graph-level prediction tasks^46^. Its advantages lie in its ability to automatically extract latent features from the graph without requiring manual feature engineering or assuming specific data distributions. Additionally, GNNs can adapt to graphs of varying scales and complexities, demonstrating strong generalization and expressive capabilities. These characteristics align well with the requirements for CDA prediction. And in capturing local depth information, subgraph structure can obtain more detailed information than node and path^47,48^. Inspired by the aforementioned article^49^, this paper proposes a recursive approach to constructing a GNN subgraph model.

As shown in Figure 8B, we use the relations between entities in the knowledge graph to recursively construct n-hop subgraph neural networks, which establish the deep information transmission between nodes in the knowledge graph. To create a subgraph for a triplet (*d, d* − *circ, circ*) in the knowledge graph, three steps are generally involved: first, we extract the neighborhood information of nodes *d* and *circ* separately; then, we take out the intersection points of these two neighborhoods to construct a subgraph; finally, we perform message propagation and subgraph encoding on the subgraph. How to obtain the two neighborhoods of *d* and *circ* becomes the key to the problem. According to the characteristics of node connectivity in the knowledge graph, we transform the problem into: recursively constructing an n-hop subgraph neural network for a node *d*, using the recursive method, we can effectively utilize the shared edges and nodes in the knowledge graph.

We denote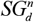 as the n-hop subgraph neural network for node *d*, 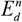 as the edge set of 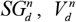 as the node set of 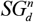, and introduce a new node set 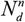, which represents the set of nodes that are not in 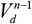 and are in 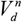, i.e. 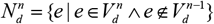. The construction method of n-hop subgraph neural network for a node *d* is as follows: First, we create a 1-hop subgraph neural network 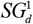 for node *d*. From the analysis of the knowledge graph, 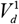is the set of nodes that are directly connected to *d*, 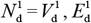 is the set of edges that are directly connected to *d*. Then, for the n-hop subgraph neural network 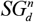for node *d*, we use 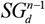 to create it.

The n-hop subgraph neural network 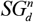 are as follows:

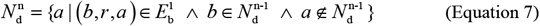

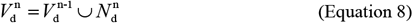

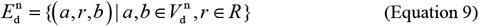

According to the above three recursive formulas, we can obtain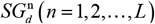, where *L* is the number of layers of the graph neural network, which reflects the depth of local information mining for triplet data by the model. Through the recursive method, we use the relationship between nodes to establish a subgraph neural network for each node, which is easier to capture local depth information^50^.

### C. Attention Aggregation to Obtain Prediction Scores

On top of the graph neural network, we use the attention layer aggregation method to update the weights of each node and each edge in the subgraph neural network, thereby selecting the strong association information in the graph neural network^51,52^. The existence of strong association information between two nodes in the knowledge graph means that the relationship between the two entities is closer, which reflects that our CDA prediction model is interpretable.

In our model, the structure that extracts the local features of the graph and fully utilizes the surrounding node information is multiple attention layers. We use the attention mechanism to assign different attention weights to different nodes and edges of the graph neural network we built. From the perspective of the model’s interpretability, the different attention weights of the attention layer control the importance and influence of different edges in the graph. The diagram of the model allocating attention weights is shown in Figure 8C.

Graph neural networks aggregate the entities and relationships of subgraphs within the messaging framework as follows:

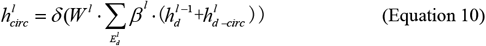

The attention weight *β*^*l*^ on edge (*d, d* − *circ, circ*) is

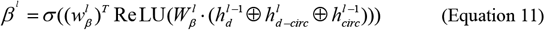

In the above formula,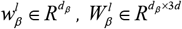,and ⊕ is the concatenation operator.

Notably, we employ the *sigmoid* function instead of *softmax attention*. This choice ensures that within the same subgraph neural network, a node can choose multiple highly correlated edges. Specifically, in the context of CDA prediction, this structure allows an entity to have relationships with multiple other entities, aligning with scenarios involving potential CDA associations. After the attention aggregation of L-layer neural networks,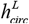 can be used to construct a scoring function for the model, so we set the scoring function to the following form:

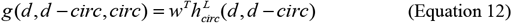

With *w* ∈ *R*^*d*^ .Consequently, we employ a multi-class logarithmic loss function as the scoring metric for our model^53^. The loss function formula is shown in formula 13:

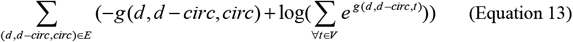

The first part of the loss function is positive triples (*d, d* − *circ, circ*) in KG,and the second part contains the scores of all triples with the same head entity and relation (*d, d* − *circ, e*) ^54^. The training parameters of the model 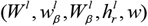 are randomly initialized and optimized using the stochastic gradient descent algorithm^30^.

## SUPPLEMENTAL INFORMATION

Supplementary data are available online.

## AUTHOR CONTRIBUTIONS

M.M., Y.W. and L.F. conceived the project, conducted the experiments, analyzed the results, and wrote the manuscript. Y.X., Q.P., H.L.and H.S. reviewed the manuscript. M.M. and Y.W. contributed to the work equllly.

## ACKNOWLEDGMENTS

This research was supported by Zhejiang Provincial Natural Science Foundation of China under Grant No. LQ23F020018, Natural Science Basic Research Program of Shaanxi (Program No. 2023-JC-QN-0737), Sichuan Science and Technology Program (No.2023NSFSC1416), National Natural Science Foundation of China (Grant No. 62303372), Young Talent Fund of Xi’an Association for Science and Technology (No. 959202313033), Project funded by China Postdoctoral Science Foundation(No. 2023M742794), and Postdoctoral Research Project in Shaanxi Province.

At the same time, we would like to thank Jinyang Wu for his support and help in this research.

## DECLARATION OF INTERESTS

The authors declare no competing of interests.

## DATAAND CODE AVAILABILITY

The code and datasets are publicly available at https://github.com/maoyuanma/KGRACDA.

## Notes

https://github.com/maoyuanma/KGRACDA

### Competing Interest Statement

The authors have declared no competing interest.

