## Supplemental Texts, Figures and Tables for "KGRACDA: A Model Based on Knowledge Graph from Recursion and Attention Aggregation for CircRNA-disease Association Prediction"

#### Supplementary Information of Methods

#### 1. Distribution of entities and relationships in the dataset

As shown in Figure S1-S3, we perform a more detailed visualization of the elements and relationships distribution of the three datasets. As can be seen from Figure S1-S3, whether it is the dataset1 and dataset3 with small data volume, or the heterogeneous dataset2 with many elements and relationships, most entities still have a degree of single digits, indicating that this is a very sparse knowledge graph, and the distribution of the dataset conforms to the distribution of the reality, and our experiments on the three datasets have credibility and persuasiveness.

#### 2. Performance comparison

In this section, we introduce some experimental details. As shown in Table S1 and Table S2, we list the detailed information of the performance comparison between KGRACDA and other 9 CDA prediction models on dataset2 and dataset3, where KGRACDA has the best comprehensive performance on dataset2 and dataset3. In addition, Table S3 shows the statistical significance of the difference between KGRACDA and other 9 methods, which further demonstrates the effectiveness of KGRACDA in accurately identifying the potential associations between circRNA and disease.

#### 3. Ablation experiments

The results of the ablation experiments of KGRACDA and other eight different knowledge graph models on dataset2 and dataset3 are shown in Figure S4-S5, which indicate that KGRACDA has the best comprehensive performance on these two datasets. The detailed information of the performance comparison between KGRACDA and other eight knowledge graph models is shown in Table S4-S5. Similarly, Table S6 shows the statistical significance of the difference between KGETCDA and other eight knowledge graph methods, reflecting the superiority of KGRACDA in predicting CDA. In addition, as shown in Figure S6-S7, we studied the impact of the number of GNN layers and the dimension of attention layer on the model prediction results on dataset2 and dataset3, and the evaluation metrics include AUC, AUPR and time consumption.

### Figures

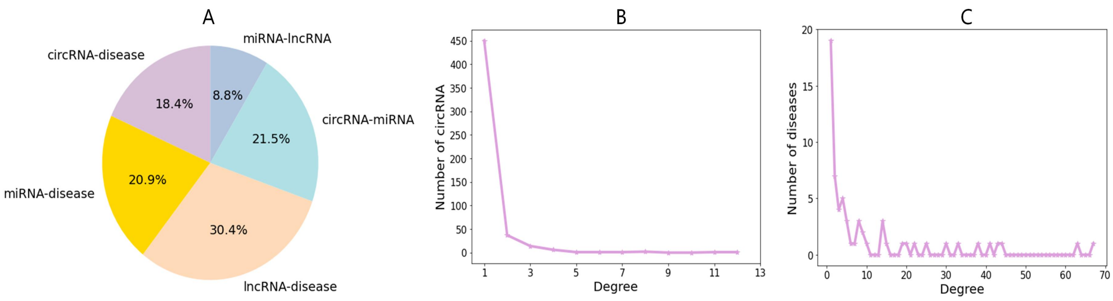


**Figure S1.** The visualization of dataset 1. **(A)** Visualize the proportion of each relationship in the dataset 1 by pie chart. **(B)** The degree distribution of dataset 1. It represents the number of corresponding circRNA associating with different numbers of disease in dataset 1. **(C)** The degree distribution of dataset 1. It represents the number of corresponding diseases associating with different numbers of circRNA in dataset1.

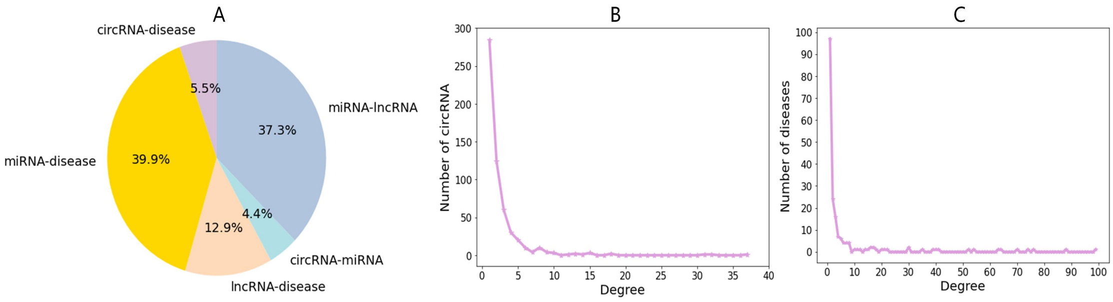


**Figure S2.** The visualization of dataset 2. **(A)** Visualize the proportion of each relationship in the dataset 2 by pie chart. **(B)** The degree distribution of dataset 2. It represents the number of corresponding circRNA associating with different numbers of disease in dataset 2. **(C)** The degree distribution of dataset 2. It represents the number of corresponding diseases associating with different numbers of circRNA in dataset 2.

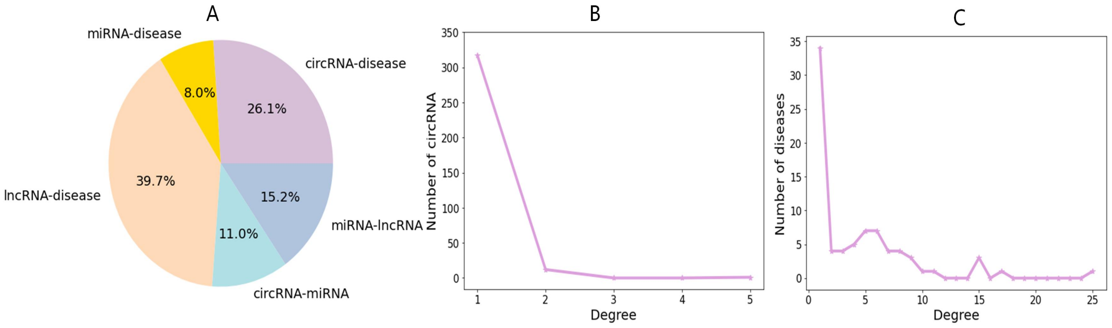

**Figure S3.** The visualization of dataset 3. **(A)** Visualize the proportion of each relationship in the dataset 3 by pie chart. **(B)** The degree distribution of dataset 3. It represents the number of corresponding circRNA associating with different numbers of disease in dataset 3. **(C)** The degree distribution of dataset 3. It represents the number of corresponding diseases associating with different numbers of circRNA in dataset 3.

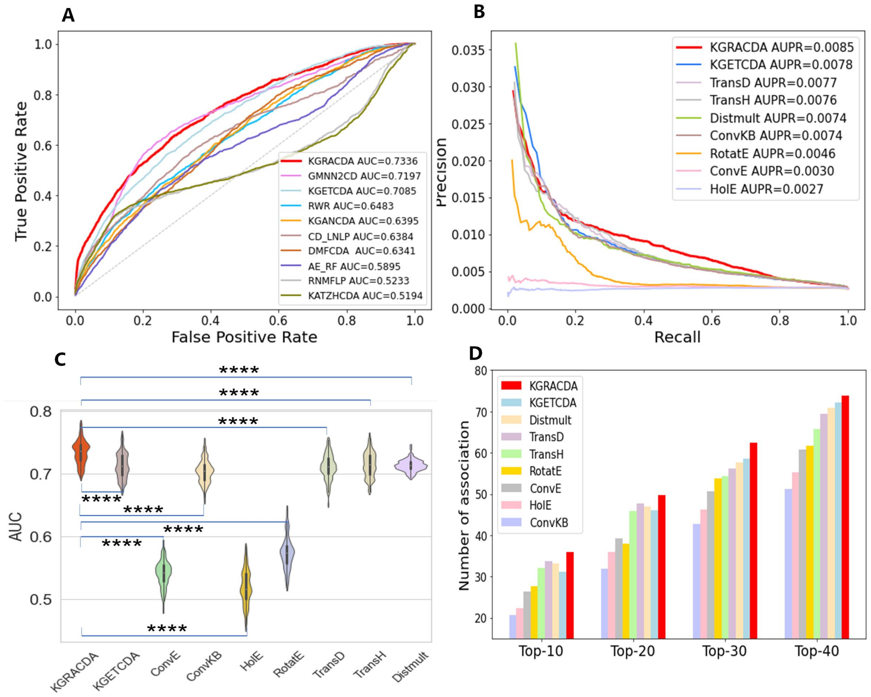


**Figure S4.** The comparison of KGRACDA with other KG methods on dataset2. **(A)** The AUC comparison **(B)** The AUPR comparison **(C)** The AUC indicators of multiple experiments are visualized in the form of violin graphs, marked with the level of significance difference for statistical hypothesis testing (**** for P-values<0.0001). **(D)** The comparison of the number of correctly identified CDA

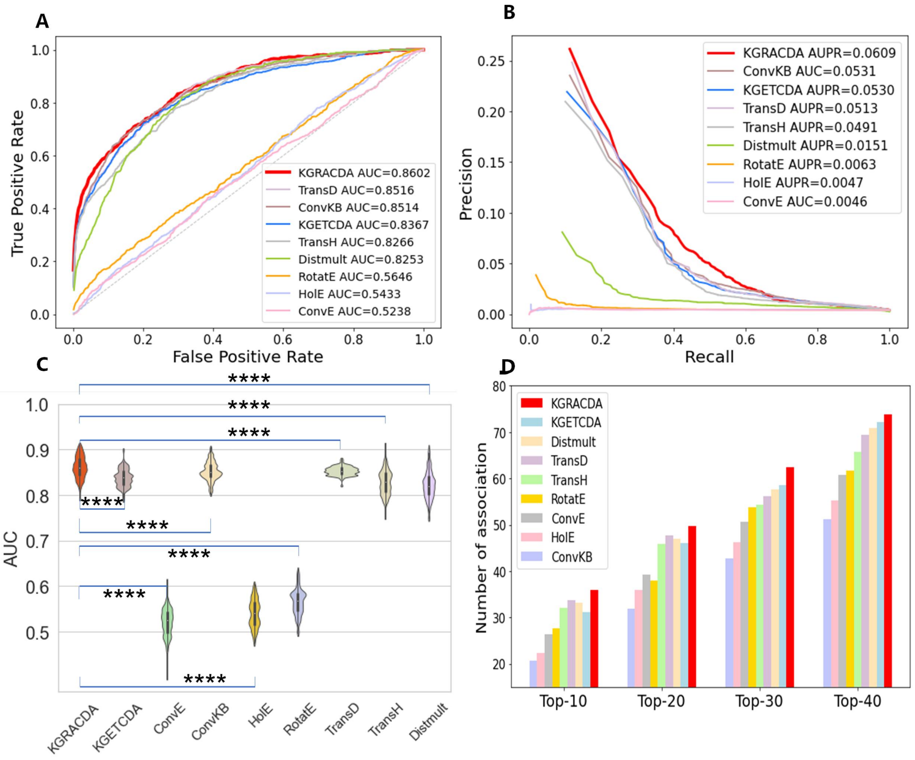

**Figure S5.** The comparison of KGRACDA with other KG methods on dataset3. **(A)** The AUC comparison (**B**) The AUPR comparison **(C)** The AUC indicators of multiple experiments are visualized in the form of violin graphs, marked with the level of significance difference for statistical hypothesis testing (**** for P-values<0.0001). **(D)** The comparison of the number of correctly identified CDA

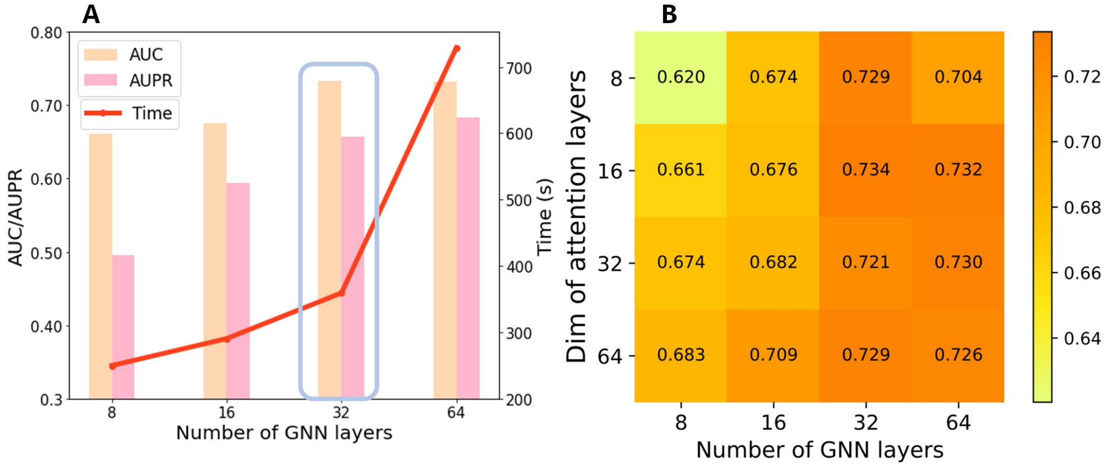


**Figure S6.** The results of ablation experiments on dataset2. **(A)** The effect of different GNN layers dimensionality. We select 32 here after considering both AUC and AUPR performance, and time consumption. For better visualization, AUPR is uniformly stretched to better highlight the trend of increase and decrease. (**B**) The effect of the number of GNN layers and the dim of attention layers.

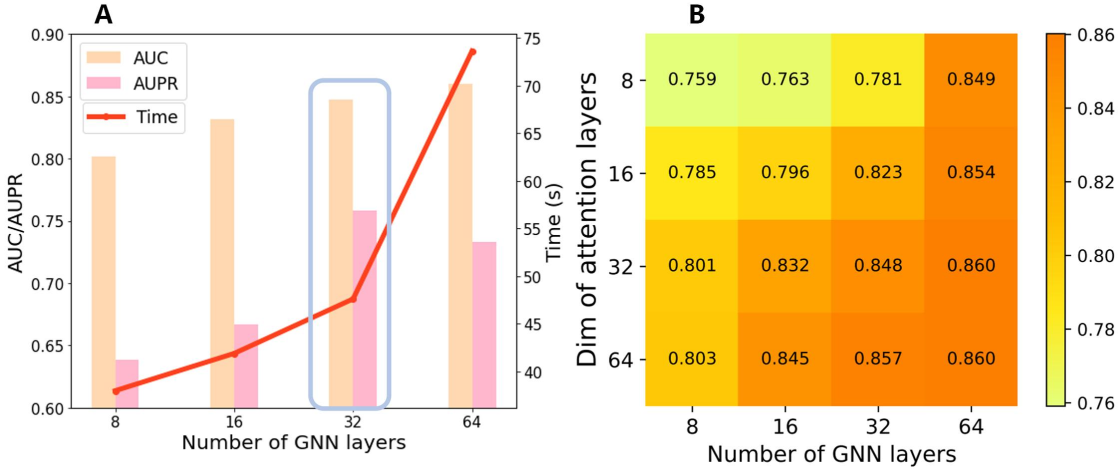


**Figure S7.** The results of ablation experiments on dataset3. **(A)** The effect of different GNN layers dimensionality. We select 32 here after considering both AUC and AUPR performance, and time consumption. For better visualization, AUPR is uniformly stretched to better highlight the trend of increase and decrease. (**B**) The effect of the number of GNN layers and the dim of attention layers.

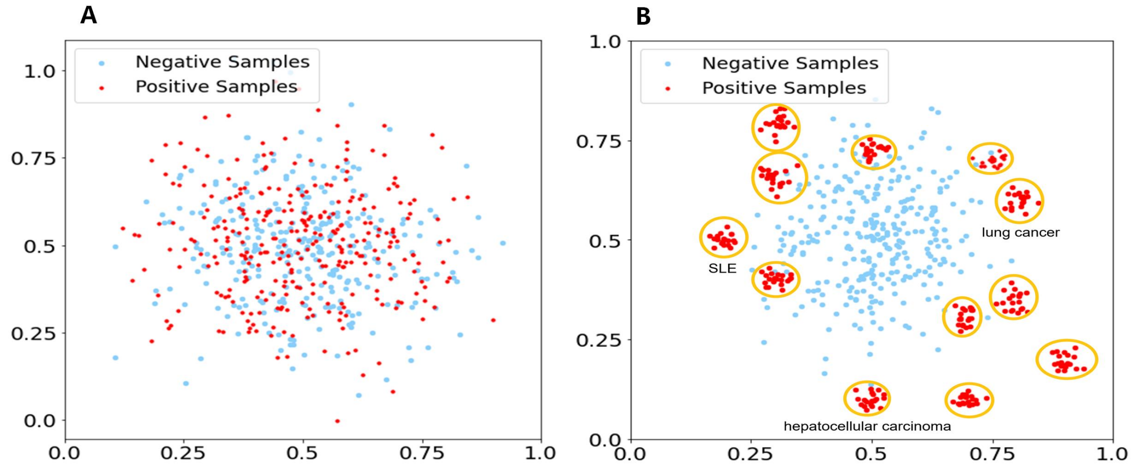


**Figure S8.** The visualization of the feature expression process using PCA on the dataset2. (A) The initial two-dimensional representation. (B) The representation after processing by KGRACDA.

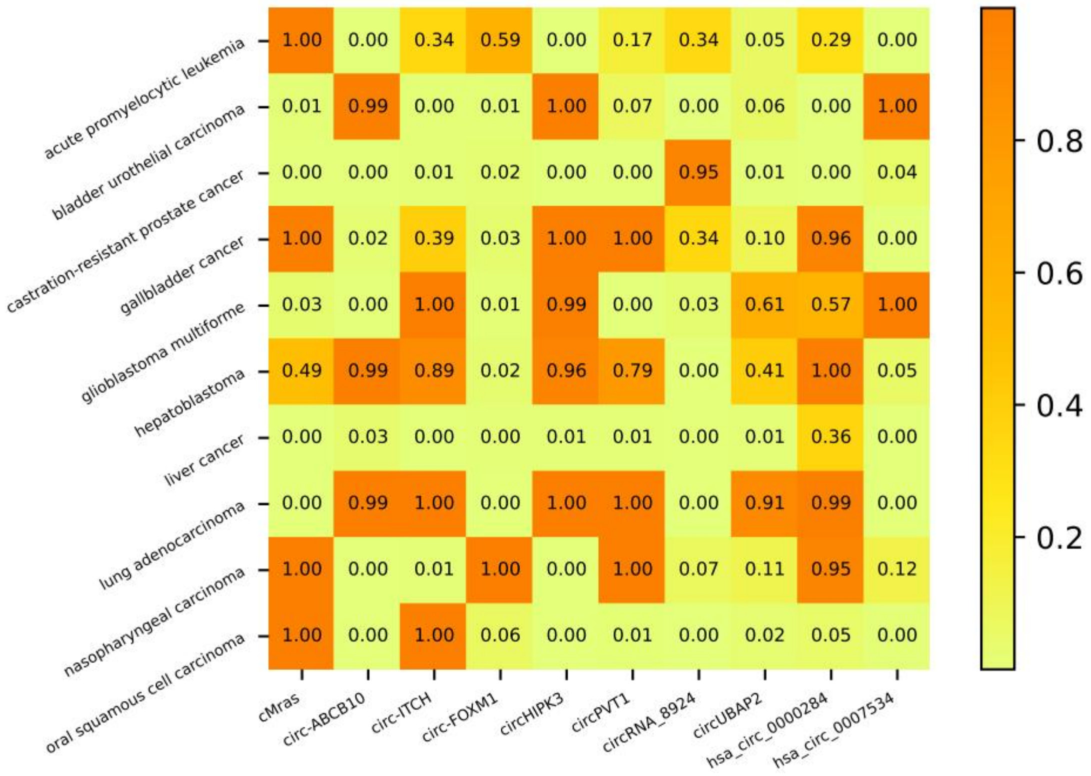


**Figure S9.** The visualization of the score matrix of randomly selected 10 circRNAs and diseases after passing the KGRACDA model.

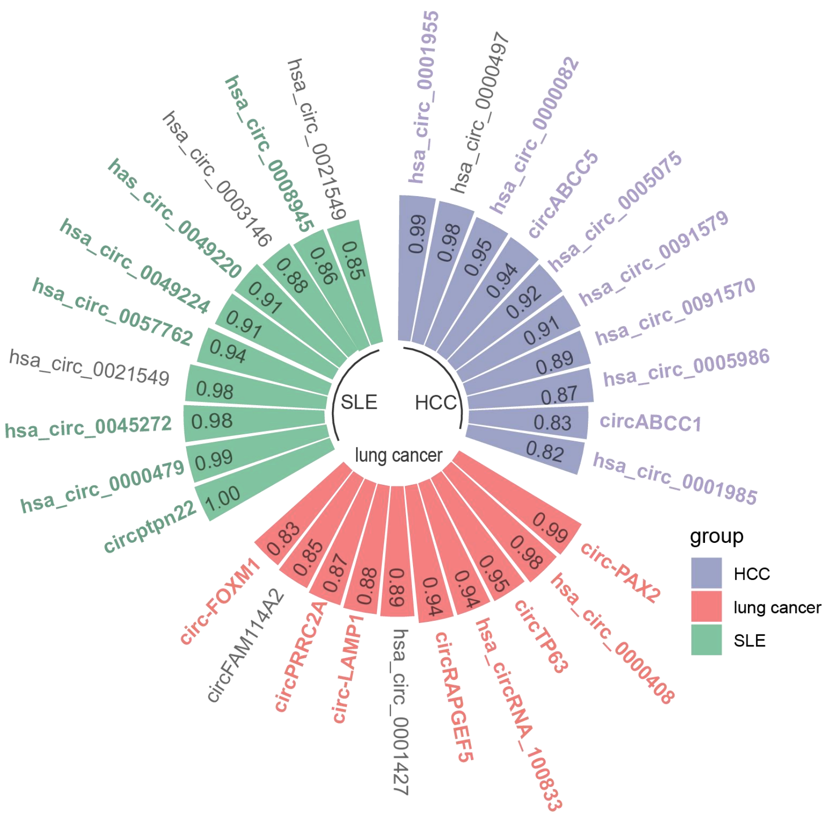


**Figure S10.** The visualization of the model case studies. The circRNA with color and bold font in the figure is the circRNA with literature support, while the circRNA with black font is the circRNA without literature support at present.

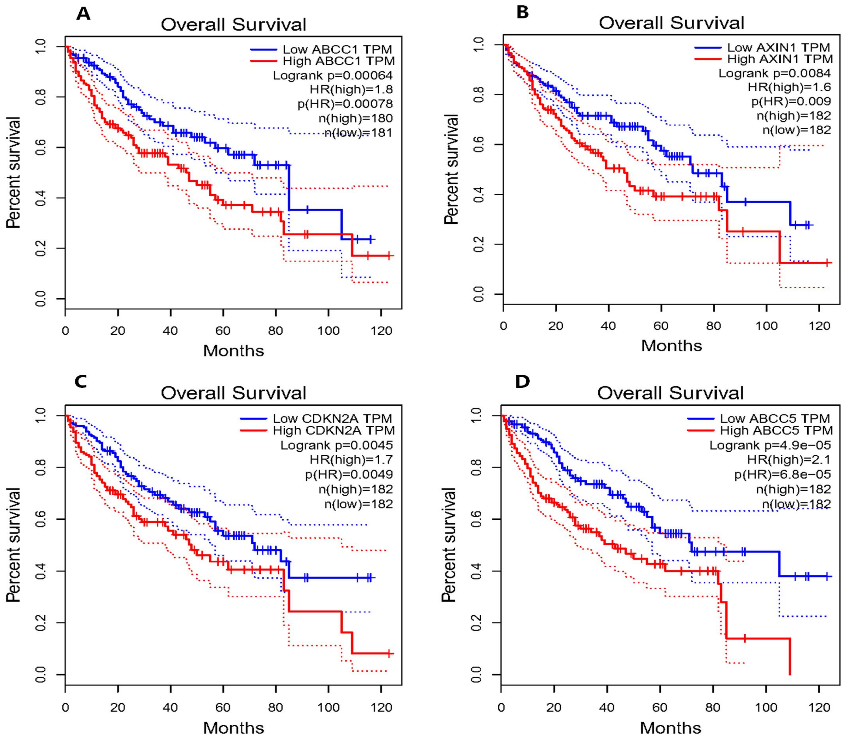


**Figure S11.** The survival curves for high and low risk patients for hepatocellular carcinoma datasets in TCGA. **(A)** ABCC1 for circABCC1. **(B)** AXIN1 for hsa_circ_0001985. **(C)** CDKN2A for hsa_circ_0000082. **(D)** ABCC5 for circABCC5.

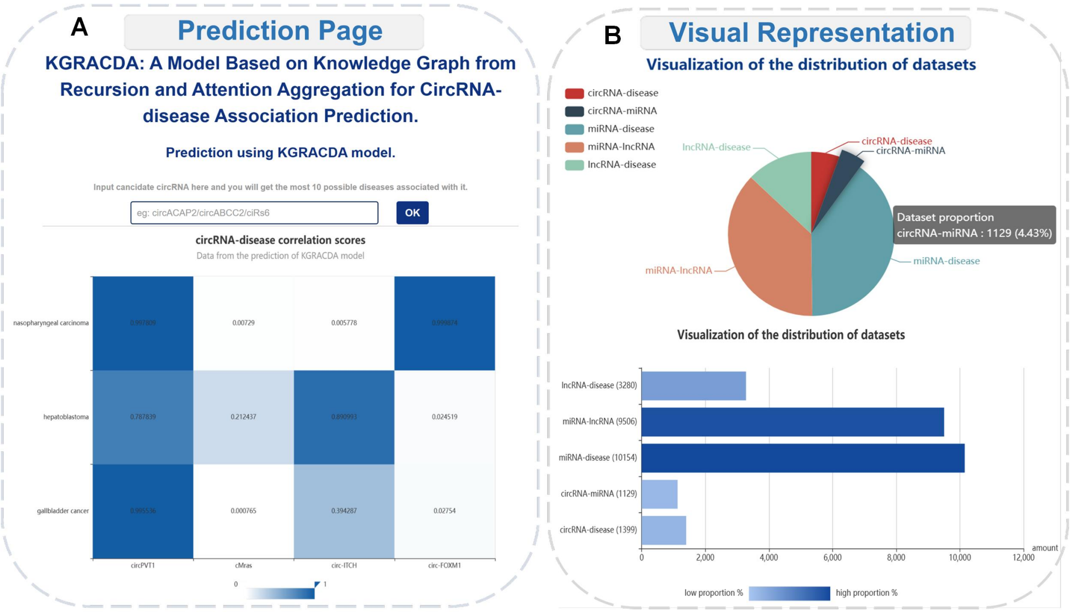


**Figure S12.** Introduction to the new features of HNRBase v2.0. (A) Many-to-many predictive heat map (B) Visual presentation of dataset information

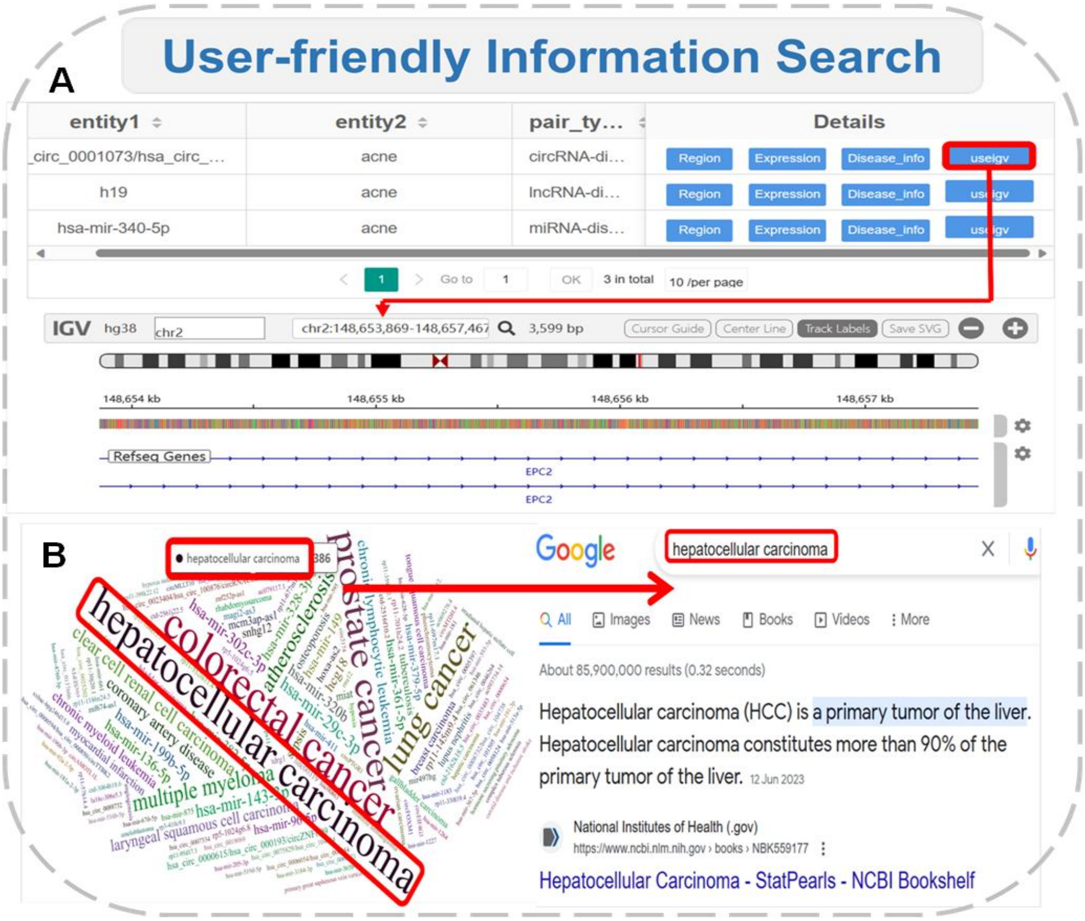


**Figure S13.** HNRBase v2.0 provides new search services for users. (A) The igv web embedding tool is used to provide circRNA corresponding genome information. (B) Examples of circRNA and disease jump search engine

### Tables

**Table S1**. The detailed information of the results of ten models on three datasets

| Dataset | Methods | Acc | Pre | Recall | F1 |
|---|---|---|---|---|---|
| Dataset1 | **KGRACDA** | **0.4988** | **0.9291** | **0.0056** | **0.0111** |
| | KGETCDA | 0.4955 | 0.9128 | 0.0089 | 0.0163 |
| | KGANCDA (2022) | 0.4950 | 0.8278 | 0.0071 | 0.0134 |
| | GMNN2CD (2022) | 0.4953 | 0.8773 | 0.0068 | 0.0135 |
| | RNMFLP (2022) | 0.4953 | 0.8687 | 0.0066 | 0.0129 |
| | AE-RF (2021) | 0.4950 | 0.8089 | 0.0052 | 0.0103 |
| | DMFCDA (2021) | 0.4939 | 0.6096 | 0.0028 | 0.0056 |
| | CD-LNLP (2019) | 0.4948 | 0.7781 | 0.0056 | 0.0111 |
| | RWR (2018) | 0.4950 | 0.8153 | 0.0064 | 0.0124 |
| | KATZHCDA(2018) | 0.4947 | 0.7752 | 0.0061 | 0.0121 |
| Dataset2 | **KGRACDA** | **0.4956** | **0.7294** | **0.0065** | **0.0129** |
| | KGETCDA (2023) | 0.4953 | 0.7119 | 0.0051 | 0.0098 |
| | KGANCDA (2022) | 0.4949 | 0.6438 | 0.0041 | 0.0080 |
| | GMNN2CD (2022) | 0.4953 | 0.7230 | 0.0051 | 0.0101 |
| | RNMFLP (2022) | 0.4943 | 0.5288 | 0.0039 | 0.0076 |
| | AE-RF (2021) | 0.4946 | 0.5944 | 0.0036 | 0.0071 |
| | DMFCDA (2021) | 0.4949 | 0.6284 | 0.0040 | 0.0077 |
| | CD-LNLP (2019) | 0.4949 | 0.6425 | 0.0045 | 0.0086 |
| | RWR (2018) | 0.4948 | 0.6446 | 0.0042 | 0.0082 |
| | KATZHCDA 2018) | 0.4943 | 0.5249 | 0.0039 | 0.0075 |
| Dataset3 | **KGRACDA** | **0.4978** | **0.8450** | **0.0103** | **0.0206** |
| | KGETCDA (2023) | 0.4940 | 0.8390 | 0.0129 | 0.0233 |
| | KGANCDA (2022) | 0.4949 | 0.7353 | 0.0096 | 0.0177 |
| | GMNN2CD (2022) | 0.4939 | 0.8274 | 0.0093 | 0.0179 |
| | RNMFLP (2022) | 0.4931 | 0.7306 | 0.0085 | 0.0163 |
| | AE-RF (2021) | 0.4930 | 0.6961 | 0.0062 | 0.0121 |
| | DMFCDA (2021) | 0.4919 | 0.5891 | 0.0044 | 0.0087 |
| | CD-LNLP (2019) | 0.4929 | 0.7004 | 0.0069 | 0.0134 |
| | RWR (2018) | 0.4933 | 0.7478 | 0.0079 | 0.0153 |
| | KATZHCDA(2018) | 0.4928 | 0.6985 | 0.0080 | 0.0155 |

**Table S2**. The number of correctly identified associations of all models on three datasets

| Dataset | Methods | TOP-10 | TOP-20 | TOP-30 | TOP-40 |
|---|---|---|---|---|---|
| Dataset1 | **KGRACDA** | **27.8** | **35.5** | **41.1** | **49.6** |
| | KGETCDA (2023) | 23.4 | 33.6 | 42.8 | 51.0 |
| | KGANCDA (2022) | 17.6 | 23.6 | 30.7 | 36.9 |
| | GMNN2CD (2022) | 10.4 | 20.8 | 29.6 | 37.2 |
| | RNMFLP (2022) | 10.0 | 20.4 | 28.6 | 39.4 |
| | AE-RF (2021) | 5.2 | 12.4 | 19.4 | 31.0 |
| | DMFCDA (2021) | 0.2 | 1.0 | 2.0 | 3.6 |
| | CD-LNLP (2019) | 8.8 | 15.0 | 23.0 | 27.8 |
| | RWR (2018) | 11.8 | 20.0 | 29.6 | 37.2 |
| | KATZHCDA (2018) | 10.8 | 18.6 | 27.4 | 34.6 |
| Dataset2 | **KGRACDA** | **36.0** | **49.8** | **62.4** | **73.9** |
| | KGETCDA (2023) | 31.2 | 46 | 58.6 | 72.2 |
| | KGANCDA (2022) | 20.0 | 32.9 | 42.2 | 51.9 |
| | GMNN2CD (2022) | 26 | 38.6 | 51.4 | 60.4 |
| | RNMFLP (2022) | 26.4 | 39.2 | 50.4 | 61.0 |
| | AE-RF (2021) | 11.0 | 21.8 | 31.0 | 39.0 |
| | DMFCDA (2021) | 17.8 | 28.0 | 37.4 | 45.8 |
| | CD-LNLP (2019) | 27.4 | 43.4 | 55.4 | 63.4 |
| | RWR (2018) | 20.0 | 32.2 | 41.2 | 53.2 |
| | KATZHCDA (2018) | 24.2 | 38.6 | 49.0 | 60.4 |
| Dataset3 | **KGRACDA** | **51.2** | **58.7** | **61.4** | **66.1** |
| | KGETCDA (2023) | 46.4 | 55.0 | 60.8 | 67.2 |
| | KGANCDA (2022) | 30.2 | 37.4 | 44.2 | 47.2 |
| | GMNN2CD (2022) | 9.3 | 20.0 | 32.4 | 41.2 |
| | RNMFLP (2022) | 9.6 | 21.4 | 30.6 | 41.0 |
| | AE-RF (2021) | 14.8 | 25.5 | 35.9 | 45.3 |
| | DMFCDA (2021) | 2.2 | 3.2 | 4.8 | 7.0 |
| | CD-LNLP (2019) | 4.6 | 8.4 | 15.4 | 22.4 |
| | RWR (2018) | 5.6 | 14.4 | 23.0 | 34.6 |
| | KATZHCDA (2018) | 10.0 | 20.8 | 28.8 | 40.2 |

**Table S3**. Statistical significance of differences between KGRACDA and the other nine methods

| | Dataset | KGETCDA | KGANCDA | GMNN2CD | RNMFLP | AE-RF | DMFCDA | CD-LNLP | RWR | KATZHCDA |
|---|---|---|---|---|---|---|---|---|---|---|
| P-value | Dataset1 | 1.216e-10 | 1.100e-10 | 1.100e-10 | 1.101e-10 | 1.103e-10 | 1.103e-10 | 1.095e-10 | 1.092e-10 | 1.091e-10 |
| | Dataset2 | 1.263e-10 | 1.103e-10 | 1.891e-10 | 1.098e-10 | 1.103e-10 | 1.619e-10 | 1.097e-10 | 1.096e-10 | 1.097e-10 |
| | Dataset3 | 1.237e-10 | 1.101e-10 | 1.205e-10 | 1.096e-10 | 1.102e-10 | 1.097e-10 | 1.094e-10 | 1.095e-10 | 1.099e-10 |

**Table S4**. The detailed information of the results of KGRACDA and other eight KG models on three datasets

| Dataset | Methods | recall | precision | F1 | AUC | AUPR |
|---|---|---|---|---|---|---|
| Dataset1 | KGRACDA | 0.9291 | 0.0056 | 0.0111 | 0.9312 | 0.0359 |
| | KGETCDA (2023) | 0.9128 | 0.0089 | 0.0167 | 0.9113 | 0.0260 |
| | RotatE (2019) | 0.7141 | 0.0047 | 0.0094 | 0.7100 | 0.0060 |
| | ConvE (2018) | 0.5108 | 0.0028 | 0.0057 | 0.5040 | 0.0029 |
| | ConvKB (2018) | 0.9186 | 0.0088 | 0.0166 | 0.9172 | 0.0256 |
| | HolE (2016) | 0.5229 | 0.0029 | 0.0058 | 0.5163 | 0.0029 |
| | Distmult (2015) | 0.9200 | 0.0089 | 0.0177 | 0.9185 | 0.0253 |
| | TransD (2015) | 0.9128 | 0.0085 | 0.0166 | 0.9113 | 0.0255 |
| | TransH (2014) | 0.8983 | 0.0086 | 0.0171 | 0.8966 | 0.0234 |
| Dataset2 | KGRACDA | 0.7294 | 0.0065 | 0.0129 | 0.7336 | 0.0085 |
| | KGETCDA (2023) | 0.7119 | 0.0051 | 0.0098 | 0.7085 | 0.0078 |
| | RotatE (2019) | 0.5762 | 0.0038 | 0.0075 | 0.5711 | 0.0046 |
| | ConvE (2018) | 0.7088 | 0.0030 | 0.0060 | 0.5400 | 0.0030 |
| | ConvKB (2018) | 0.7090 | 0.0049 | 0.0100 | 0.7056 | 0.0074 |
| | HolE (2016) | 0.5161 | 0.0027 | 0.0052 | 0.5102 | 0.0027 |
| | Distmult (2015) | 0.7173 | 0.0050 | 0.0101 | 0.7140 | 0.0074 |
| | TransD (2015) | 0.7140 | 0.0051 | 0.0102 | 0.7106 | 0.0077 |
| | TransH (2014) | 0.7150 | 0.0052 | 0.0103 | 0.7116 | 0.0076 |
| Dataset3 | KGRACDA | 0.8450 | 0.0103 | 0.0206 | 0.8602 | 0.0609 |
| | KGETCDA (2023) | 0.8390 | 0.0129 | 0.0233 | 0.8367 | 0.0530 |
| | RotatE (2019) | 0.5723 | 0.0056 | 0.0110 | 0.5646 | 0.0063 |
| | ConvE (2018) | 0.5323 | 0.0045 | 0.0090 | 0.5238 | 0.0046 |
| | ConvKB (2018) | 0.8434 | 0.0132 | 0.0261 | 0.8514 | 0.0531 |
| | HolE (2016) | 0.5514 | 0.0046 | 0.0092 | 0.5433 | 0.0047 |
| | Distmult (2015) | 0.8278 | 0.0047 | 0.0140 | 0.8253 | 0.0151 |

| | | | | | | |
|---|---|---|---|---|---|---|
| | TransD (2015) | 0.8436 | 0.0131 | 0.0259 | 0.8516 | 0.0513 |
| | TransH (2014) | 0.8290 | 0.0125 | 0.0246 | 0.8266 | 0.0491 |

**Table S5**. The number of correctly identified associations of KGRACDA and other eight KG models on three datasets

| Dataset | Methods | TOP-10 | TOP-20 | TOP-30 | TOP-40 |
|---|---|---|---|---|---|
| Dataset1 | KGRACDA | 27.8 | 35.5 | 41.1 | 49.6 |
| | KGETCDA (2023) | 23.4 | 33.6 | 42.8 | 51.0 |
| | RotatE (2019) | 4.3 | 10.7 | 19.8 | 29.2 |
| | ConvE (2018) | 22.5 | 30.6 | 40.0 | 45.7 |
| | ConvKB (2018) | 5.0 | 11.2 | 19.3 | 28.6 |
| | HolE (2016) | 8.6 | 15.4 | 22.9 | 32.1 |
| | Distmult (2015) | 24.2 | 32.1 | 43.8 | 47.7 |
| | TransD (2015) | 21.0 | 30.9 | 41.2 | 45.3 |
| | TransH (2014) | 23.5 | 32.7 | 40.5 | 47.2 |
| Dataset2 | KGRACDA | 36.0 | 49.8 | 62.4 | 73.9 |
| | KGETCDA (2023) | 31.2 | 46.0 | 58.6 | 72.2 |
| | RotatE (2019) | 27.6 | 38.0 | 53.8 | 61.7 |
| | ConvE (2018) | 26.4 | 39.3 | 50.6 | 60.8 |
| | ConvKB (2018) | 20.7 | 31.9 | 42.7 | 51.2 |
| | HolE (2016) | 22.4 | 36.0 | 46.2 | 55.2 |
| | Distmult (2015) | 33.1 | 47.0 | 57.6 | 70.9 |
| | TransD (2015) | 33.8 | 47.7 | 56.2 | 69.4 |
| | TransH(2014) | 32.0 | 45.8 | 54.3 | 65.8 |
| Dataset3 | KGRACDA | 51.2 | 58.7 | 61.4 | 66.1 |
| | KGETCDA (2023) | 46.4 | 55.0 | 60.8 | 67.2 |
| | RotatE (2019) | 2.0 | 4.3 | 5.1 | 6.9 |
| | ConvE (2018) | 9.9 | 21.7 | 33.4 | 42.4 |
| | ConvKB (2018) | 2.3 | 4.8 | 5.6 | 7.2 |
| | HolE (2016) | 3.3 | 5.1 | 6.7 | 8.0 |
| | Distmult (2015) | 47.2 | 54.2 | 58.7 | 66.2 |
| | TransD (2015) | 43.8 | 53.7 | 58.3 | 63.3 |
| | TransH (2014) | 39.9 | 48.1 | 56.5 | 62.8 |

**Table S6**. Statistical significance of differences between KGRACDA and the other nine KG methods

| | **Dataset** | **KGETCDA** | **ConvE** | **ConvKB** | **HolE** | **RotatE** | **TransD** | **TransH** | **Distmult** |
|---|---|---|---|---|---|---|---|---|---|
| P-value | Dataset1 | 1.216e-10 | 1.018e-10 | 1.264e-10 | 1.003e-10 | 1.130e-10 | 1.272e-10 | 1.258e-10 | 1.291e-10 |
| | Dataset2 | 1.263e-10 | 1.025e-10 | 1.149e-10 | 1.014e-10 | 1.188e-10 | 1.276e-10 | 1.364e-10 | 1.278e-10 |
| | Dataset3 | 1.237e-10 | 1.056e-10 | 1.174e-10 | 1.031e-10 | 1.040e-10 | 1.196e-10 | 1.209e-10 | 1.221e-10 |

**Table S7**. The summary of experimental settings

| **Hyperparameter** | **Our settings** |
|---|---|
| Training epochs | 50 |
| Lr | 2e-4 |
| Batch size | 128 |
| Dim of attention layer | 32 |
| Dim of hidden layer | 64 |
| GNN layers | 32 |
| Weight decay | 1e-3 |
| Dropout | 0.02 |
| Lr_decay | 0.999 |

**Table S8**. The parameter sensitivity analysis

| AUC | **Dataset** | 1e-2 | 1e-3 | **2e-4** | 1e-4 | 5e-5 | 1e-5 | 5e-6 |
|---|---|---|---|---|---|---|---|---|
| Learning Rate | Dataset1 | 0.8698 | 0.8709 | 0.9312 | 0.9165 | 0.8863 | 0.8701 | 0.8689 |
| | Dataset2 | 0.6783 | 0.6859 | 0.7336 | 0.7142 | 0.6813 | 0.6755 | 0.6638 |
| | Dataset3 | 0.7931 | 0.8077 | 0.8481 | 0.8602 | 0.8384 | 0.8241 | 0.8194 |
| | **Dataset** | 0.99 | 0.998 | **0.999** | 0.9998 | 0.9999 | 0.99998 | 0.99999 |
| Lr_decay | Dataset1 | 0.9103 | 0.9209 | 0.9312 | 0.9245 | 0.9189 | 0.9031 | 0.8977 |
| | Dataset2 | 0.6932 | 0.7265 | 0.7336 | 0.7108 | 0.7011 | 0.6841 | 0.6692 |
| | Dataset3 | 0.7935 | 0.8320 | 0.8548 | 0.8602 | 0.8401 | 0.8330 | 0.8257 |
| | **Dataset** | 1e-2 | 2e-3 | **1e-3** | 2e-4 | 1e-4 | 2e-5 | 1e-5 |
| Weight Decay | Dataset1 | 0.9288 | 0.9304 | 0.9312 | 0.9297 | 0.9216 | 0.9133 | 0.9073 |
| | Dataset2 | 0.7071 | 0.7216 | 0.7336 | 0.7158 | 0.7076 | 0.6891 | 0.6725 |
| | Dataset3 | 0.8115 | 0.8412 | 0.8534 | 0.8602 | 0.8544 | 0.8283 | 0.8229 |
